# Macro- and Micro-Structural Alterations in the Midbrain in Early Psychosis

**DOI:** 10.1101/2024.04.10.588901

**Authors:** Zicong Zhou, Kylie Jones, Elena I. Ivleva, Luis Colon-Perez

**Author notes:** Corresponding author: Luis Colon-Perez, PhD, University of North Texas Health Science Center, 3500 Camp Bowie Boulevard, Fort Worth, TX 76107.

## Abstract

**Introduction:** Early psychosis (EP) is a critical period in the course of psychotic disorders during which the brain is thought to undergo rapid and significant functional and structural changes ^1^. Growing evidence suggests that the advent of psychotic disorders is early alterations in the brain’s functional connectivity and structure, leading to aberrant neural network organization. The Human Connectome Project (HCP) is a global effort to map the human brain’s connectivity in healthy and disease populations; within HCP, there is a specific dataset that focuses on the EP subjects (i.e., those within five years of the initial psychotic episode) (HCP-EP), which is the focus of our study. Given the critically important role of the midbrain function and structure in psychotic disorders (cite), and EP in particular (cite), we specifically focused on the midbrain macro- and micro-structural alterations and their association with clinical outcomes in HCP-EP.

**Methods:** We examined macro- and micro-structural brain alterations in the HCP-EP sample (n=179: EP, n=123, Controls, n=56) as well as their associations with behavioral measures (i.e., symptoms severity) using a stepwise approach, incorporating a multimodal MRI analysis procedure. First, Deformation Based Morphometry (DBM) was carried out on the whole brain 3 Tesla T1w images to examine gross brain anatomy (i.e., seed-based and voxel-based volumes). Second, we extracted Fractional Anisotropy (FA), Axial Diffusivity (AD), and Mean Diffusivity (MD) indices from the Diffusion Tensor Imaging (DTI) data; a midbrain mask was created based on FreeSurfer v.6.0 atlas. Third, we employed Tract-Based Spatial Statistics (TBSS) to determine microstructural alterations in white matter tracts within the midbrain and broader regions. Finally, we conducted correlation analyses to examine associations between the DBM-, DTI- and TBSS-based outcomes and the Positive and Negative Syndrome Scale (PANSS) scores.

**Results:** DBM analysis showed alterations in the hippocampus, midbrain, and caudate/putamen. A DTI voxel-based analysis shows midbrain reductions in FA and AD and increases in MD; meanwhile, the hippocampus shows an increase in FA and a decrease in AD and MD. Several key brain regions also show alterations in DTI indices (e.g., insula, caudate, prefrontal cortex). A seed-based analysis centered around a midbrain region of interest obtained from freesurfer segmentation confirms the voxel-based analysis of DTI indices. TBSS successfully captured structural differences within the midbrain and complementary alterations in other main white matter tracts, such as the corticospinal tract and cingulum, suggesting early altered brain connectivity in EP. Correlations between these quantities in the EP group and behavioral scores (i.e., PANSS and CAINS tests) were explored. It was found that midbrain volume noticeably correlates with the Cognitive score of PA and all DTI metrics. FA correlates with the several dimensions of the PANSS, while AD and MD do not show many associations with PANSS or CAINS.

**Conclusions:** Our findings contribute to understanding the midbrain-focused circuitry involvement in EP and complimentary alteration in EP. Our work provides a path for future investigations to inform specific brain-based biomarkers of EP and their relationships to clinical manifestations of the psychosis course.

## Introduction

Early psychosis (EP) represents a critical period in the course of psychotic disorders during which early detection and comprehensive interventions substantially improve symptom and functional outcomes and have the potential to modify the subsequent disease course ^2–4^. While the clinical manifestations of EP have been extensively characterized ^5^, the development of objective, measurable biomarkers that can augment clinical diagnoses and inform actionable targets for interventions remains limited. A meta-analysis suggests that about 6% of individuals presenting with first-episode psychosis demonstrate clinically relevant findings on MRI ^6^, highlighting the significance of identifying novel MRI biomarkers that will aid in the differential diagnosis of various EP syndromes. However, once the neurological and medical causes of EP have been ruled out, clinical MRI, in its current stand, is largely non-contributary to further diagnostic algorithms, monitoring disease course and outcomes, and guiding choices of intervention. Developing refined, diagnosis-specific, yet cost-efficient and practical MRI-based biomarkers for psychiatric disorders is a high priority for the field. Comprehensive, in-depth *in vivo* brain characterization (e.g., via multimodal MRI approaches) with robust cognitive, symptom, and functional benchmarks may improve clinical prediction in EP and inform more individualized precision treatments ^7^.

Potential early biomarkers of psychotic disorders may lie in alterations in the structure, function, and connectivity of brain networks and circuits^8–10^. Network-based studies show that white matter (i.e., network edges) serves as a conduit of pathology that spreads in psychotic disorders and evolves over the disease stages ^11^. The anterior hippocampus is considered a significant hub of early functional^12^ and structural alterations in EP, with a subsequent proliferation to the posterior and prefrontal cortex.^11^ Clinical high-risk individuals who subsequently converter to psychosis display reductions in cortical thickness in the frontal cortical system ^13^. Recent large-scale machine learning approaches detected two anatomical network-based subtypes within the EP population: one starting with volumetric reductions in the hippocampus and related subcortical regions (“early subcortical” subtype), and second, with the initial alterations in insula and Broca’s area cortex (“early cortical” subtype). In both subtypes, structural alterations propagated to the rest of the brain throughout psychotic illness ^14^. Significantly, these structural MRI-based disease subtypes predicted clinical outcomes, i.e., substantially higher psychosis, depression, and anxiety, and general psychopathology scores found in the “early cortical” subtype ^14^. These findings highlight the importance of the circuit-focused MRI approaches in EP and subsequent illness course and their potential utility for clinical implication.

The midbrain is a particular brain region vulnerable to psychotic disorders. Individuals at ultra-high risk for psychosis showed increased resting cerebral blood flow in the midbrain and hippocampus ^15^. Increased regional activation in the midbrain in response to neutral stimuli has been suggested as a functional MRI biomarker of delusions in schizophrenia ^16^. Individuals with schizophrenia also show reduced functional connectivity in the hippocampus-midbrain-striatum network ^17^. In addition, PET studies in individuals with schizophrenia demonstrate elevated dopamine synthesis capacity in the substantia nigra, another hub region of the midbrain, which in turn is associated with symptom severity in schizophrenia.^18^ Midbrain involvement in mechanisms of psychosis is further supported by observations in animal models where midbrain dopaminergic signaling is altered due to efferent from the medial temporal lobe ^19^. Dopamine synthesis and signaling alterations in the midbrain and striatum are downstream from the “primary” Glu/GABA alterations in the upstream limbic system.^20,21^ In addition to the alterations in the midbrain dopamine system, other regional markers have been implicated in schizophrenia, e.g., serotonin hyperactivity^22^ and alterations in neuroinflammatory signaling via increases in IL6 and TNFα^23^.

The midbrain is one of the critical brain systems affected in EP;^24^ however, few studies have focused on the midbrain and EP. The ventral tegmental area (VTA), a critical brain region for reward learning and memory located in midbrain, is impaired in the early stages of psychosis: VTA-hippocampal functional connectivity is increased in EP, while VTA-striatal connectivity is reduced ^25,26^, suggesting differential effects of dopaminergic circuits mediating seemingly contradictory effects in behavior (i.e., hypodopaminergic like anhedonia vs hyperdopaminergic like delusion or paranoia). VTA-hippocampal functional connectivity correlates with individual differences in the overall symptom severity as determined by the Brief Psychiatric Rating Scale ^25^. More studies are needed to elucidate the role of midbrain function and structure in EP.

In this study, we used publicly available data from the HCP-EP, which aimed to generate a high-quality, comprehensive dataset in individuals with early-course psychosis (that is, within five years of psychosis onset). We focused on the data’s whole brain structural and diffusion MRI components to identify macro and micro-structural alterations in EP and their associations with clinical symptoms. We employed a stepwise approach incorporating 1) whole-brain Voxel-Based Morphometry (VBM) and Deformation-Based Morphometry (DBM) to examine regional alterations in gray matter volume; 2) a voxel-based over the entire brain and seed-based (midbrain segmentation) analyses of DTI indices capturing alterations in white matter microstructure; and 3) Tract-Based Spatial Statistics (TBSS) to examine white matter tract alterations, in EP. Following the above whole-brain analyses and given the vital role of the midbrain in EP pathophysiology (as reviewed above), we further quantified its structural properties using a seed-based approach. Finally, we performed correlation analyses to examine associations between several midbrain-centered MRI metrics and symptom dimensions of psychosis in the HCP-EP dataset.

## Materials and Methods

### Participants

The HCP-EP dataset comprises 251 subjects (183 EP and 68 matched healthy controls, Release 1.1). We included subjects with T1w and DTI data (123 EP and 56 controls). The EP cohort consisted of subjects within five years of psychosis onset, including affective and non-affective psychoses. Inclusionary diagnoses were DSM-5^27^ diagnosis of schizophreniform disorder, schizophrenia, schizoaffective disorder, psychosis unspecified type, delusional disorder, or brief psychotic disorder, major depressive disorder with psychotic features (single and recurrent episodes), or bipolar disorder with psychotic features (including most recent episode depressed and manic types), all with onset within five years before study entry. We obtained approval from NIH/NIMH through a Data Use Certification agreement (OMB Control Number: 0925-0667).

### MRI

This study focused on the 1) T1w MPRAGE (0.8mm isotropic resolution) and the 2) diffusion-weighted MRI sequence (MB acceleration factor of 4, 92 directions in each shell (b=1500 and 3000) acquired twice: once with AP and once with PA phase encoding). More details on the data acquisition parameters can be found on the HCP-EP website. (https://www.humanconnectome.org/storage/app/media/documentation/HCP-EP1.1/Appendix_1_HCP-EP_Release_Imaging_Protocols.pdf)

### Voxel-Based Morphometry (VBM) and Deformation-Based Morphometry (DBM)

We employed Anatomical Normalization Tools (ANTs) for the whole brain T1 structural analysis. We used 123 EP subjects and 53 controls with complete T1w images that passed quality control procedures (detailed in Supplementary Methods). To examine whether results are reproducible over distinct template spaces, the VBM ^28^ procedure was executed thrice using three distinct structural atlases (2 general atlases and one local template): (1) the common standard template MNI152, (2) OASIS ^29^ and (3) a local template constructed from 56 healthy controls (ANTs’ antsMultivariateTemplateConstruction2). Each T1w image was registered (ANTs’ antsRegistrionSyN) onto each template, for VBM^30^. In addition, we employed DBM to capture the spatial variability of shape changes among the deformation fields from image registration ^31^. DBM measures variabilities of the registration movements using the mathematical quantity Jacobian determinant (JD). In contrast, VBM measures the variabilities in image intensity differences and is considered to somewhat overlook spatial shape changes during linear and non-linear image registration steps. DBM performs a similar statistical analysis to VBM but considers the values of JD, whose value ranges around 1, indicating regional shrinkage for values less than one and expansion for values greater than 1.

### DTI and Tract-Based Spatial Statistics (TBSS)

Diffusion data was motion corrected (FSL’s eddy), skull stripped (BET), and corrected for field inhomogeneities (FSL’s topup). We generated DTI indices (FSL’s dtifit): fractional anisotropy (FA), mean diffusivity (MD), and axial diffusivity (AD). Then, the data was registered to the MNI atlas and analyzed for group differences at the voxel level by t-tests, similar to VBM. TBSS was performed based on FAs following the FSL’s TBSS guidelines (FSL’s randomize, 123 EP subjects and 56 controls with complete and adequate DW images) ^32^.

### Midbrain-focused analyses

Given our focus on the midbrain, we segmented the brainstem region-of-interest (ROI) using the fressurfer and manually segmented portions of the pons and medulla. We used the first slice where there was a clear distinction of the pons as a marker of the inferior portion of the midbrain. This segmented midbrain structure was then registered to all DTI scalar maps for seed-based analysis of DTI indices.

### Symptom data

The Positive and Negative Syndrome Scale (PANSS) ^33^ and the Clinical Assessment Interview for Negative Symptoms (CAINS)^34^ were obtained as part of the behavioral dataset of the HCP-EP. Both scales are well-validated, reliable, and extensively utilized in the assessment of current symptom severity in psychosis populations, including EP. Each section of the PANSS test displays an internal reliability of 0.73, 0.83, and 0.79 for positive, negative, and general psychopathology scales, respectively, and interrater reliability of r = 0.83, 0.85, 0.87 for positive, negative, and general psychopathology scales, respectively;^35^ meanwhile the CAINS has an internal reliability of 0.76 and test-retest reliability of r = 0.69.^34^

### Statistical Analyses

VBM and DBM statistics were completed in Python using Numpy and Scipy packages with voxel-wise t-tests (alpha = 0.05) followed by Family-Wise Error (FWE) rate correction using the False Discovery Rate. FA, AD, and MD map voxel-level t-tests (alpha = 0.05) were completed using the same Python-based approach. Seed-based midbrain outcomes were analyzed using Welch Two Sample t-tests on the mean FA, AD, and MD maps in R with the t-test function and shown with a violin plot using the ggplot function. The TBSS was carried out on FA maps, in which the general linear model steps were done using the FSL randomise function with 1000 permutations. The correlation analyses examined associations between the symptom scales (PANSS and CAINS), volume measures (i.e., whole brain JD, midbrain volumes), and DTI indices in the EP group in R using paris.panels from the package PerformanceAnalytics.

## Results

The total dataset consisted of 179 subjects (EP and controls), with 68.7% being males and 31.3% females, evenly distributed in EP (males = 61%, females = 37.4%) and controls (males = 64.3%, females = 35.7%) subsets (**Table 1**). Subjects were primarily right-handed (EP = 85.4%, controls = 80.4%). The EP cohort included 75.6% and 24.4% non-affective and affective psychosis subjects, respectively.

**Table 1.** Demographic characteristics of the HCP-EP dataset. One EP subject did not have demographic data reported.

|  | EP, n = 123 | Controls, n =56 |
| --- | --- | --- |
| Age, years, Mean (SD) | 23.06 (3.64) | 24.43 (4.27) |
| Sex, n (%) | M: 75 (61.0 %)<br>F: 46 (37.4 %)<br>N/A: 2 (1.6%) | M: 36 (64.3 %)<br>F: 20 (35.7 %) |
| Handedness, n (%) | Right: 105 (85.4 %)<br>Left: 10 (8.1 %)<br>Ambidextrous: 5 (4.1 %)<br>N/A: 3 (2.4 %) | Right: 45 (80.4 %)<br>L: 10 (17.8 %)<br>Ambidextrous: 1 (1.8 %) |
| Psychosis Phenotype, n (%) | Affective: 28 (24.4 %)<br>Non-Affective: 93 (75.6 %) | NA |
| Lifetime Antipsychotic medication exposure: months, Mean (SD) | Yes: 97 (78.9 %)<br>No: 26 (21.1 %)<br>14.06 (15.58) | NA |
| Race/Ethnicity, n (%) | Asian: 9 (7.3 %)<br>Black: 45 (36.5 %)<br>Hispanic/Latino: 4 (3.2 %)<br>Multi-racial: 7 (5.6 %)<br>White: 58 (47.1 %)<br>N/A: 2 (1.6 %) | Asian: 7 (12.5 %)<br>Black: 5 (8.9 %)<br>Multi-racial: 9 (16.1 %)<br>White: 34 (60.7 %)<br>N/A: 1 (1.8 %) |
| Education (in degrees and certificates | 6.32 (2.10) | 8.89 (2.43) |
| earned) |  |  |
| IQ<br>Standardized<br>Score, Mean<br>(SD) | 101.68 (16.95) | 115.28 (11.01) |

The DBM method is agnostic to tissue type and relies on image contrast for displacements in the registration; therefore, DBM does not suffer from tissue misclassification as VBM, and is potentially more sensitive than VBM to subtle alterations comparing healthy and patient populations.^36^ DBM revealed significant between-group morphological alterations, indicating higher displacement (i.e., JD values) in frontal regions and the superior regions within the midbrain in EP vs. healthy controls (**Fig. 1, Supplementary Figs. 4-6**). In contrast, the standard VBM yielded no significant between-group differences (**Supplementary Figs. 1-3**). All three templates produced similar patterns, indicating the reproducibility of volumetric alterations in EP vs. controls and highlighting the same regional alterations throughout the brain independent of the segmentation (i.e., tissue classification) template. In addition, we could show DBM alterations in the anterior hippocampus, frontal cortex, insula, and putamen (Fig 1).

**Figure 1.**
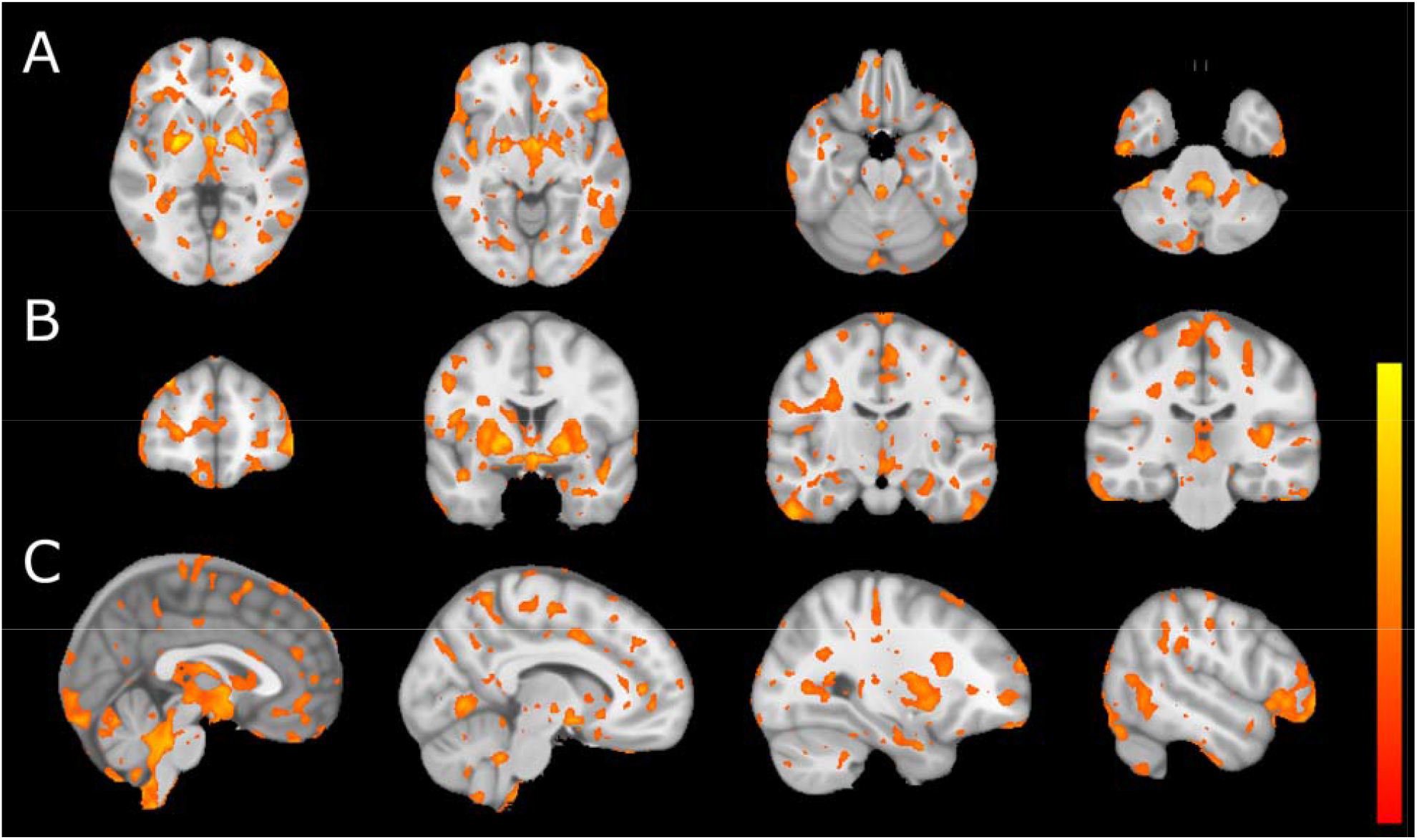
Volumetric differences in the EP vs. healthy control groups. Volumetric differences are estimated from the DBM parameter using the JD results. Yellow represents voxel-wise volume increases, and red represents reductions in EP vs. healthy controls (t = 2, q < 0.05).

To determine alterations in white matter microstructure between EP and controls, we obtained DTI indices: whole-brain maps of FA (**Supplementary Figs. 13-14**), AD (**Supplementary Figs. 15-16**), and MD (**Supplementary Figs. 17-18**). Susceptibility artifacts were modeled by AP and PA acquisition and corrected to improve image quality. FA maps registered to the MNI152 template revealed significant between-group differences in several white matter tracts (corpus callosum, cingulum, and cortico-spinal tract), with lower FA in EP vs. controls (**Fig 2**). The voxel-based analysis shows alterations in voxels in the midbrain; significantly lower FA indices were seen at the cerebral peduncles and regions adjacent to VTA in EP vs. controls, as shown in **Fig 2**. MD maps showed substantially higher (e.g., in the corpus callosum, midbrain, and basal ganglia regions) and lower (in the medial temporal lobe) MD in EP vs. controls. The MD alterations in the midbrain were localized to the VTA area (**Fig 3**). The AD maps displayed between-group differences mainly in the ventricular and peripheral CSF spaces, with lower AD in EP vs. controls (**Fig 4**). Notably, the AD results also indicated alterations in the midbrain regions (i.e., in peduncles and near VTA), similar to those captured in FA maps.

**Figure 2.**
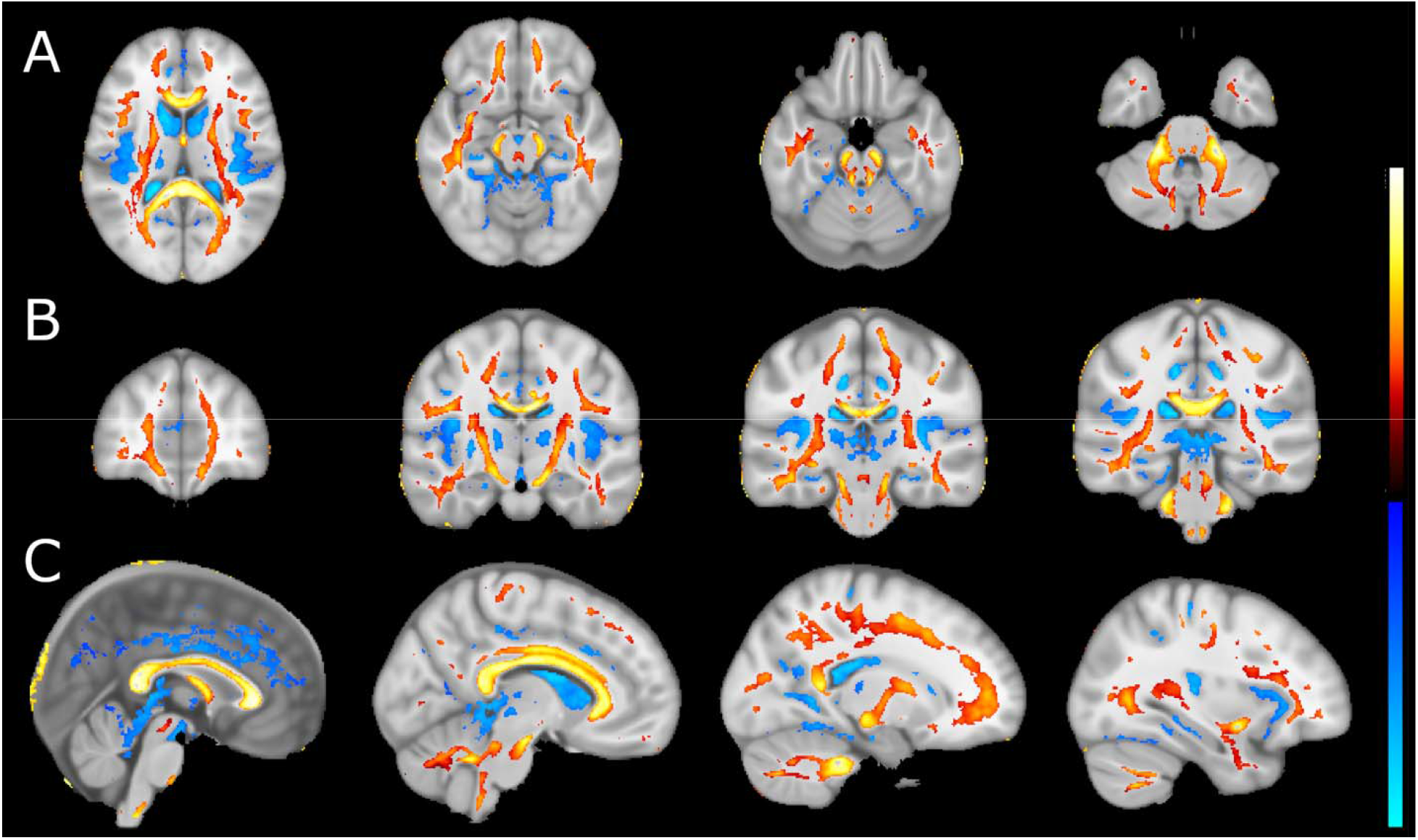
Fractional anisotropy differences in the EP vs. healthy control groups. (A) Axial, (B) coronal, and (C) sagittal sections of FA alterations between EP and controls. Differences are estimated from the FA maps registered to MNI space and following a t-test. Hot colors represent reductions in FA, while cool colors refer to increases in FA between healthy controls and EP cohorts larger than t = 6 and q < 0.05.

**Figure 3.**
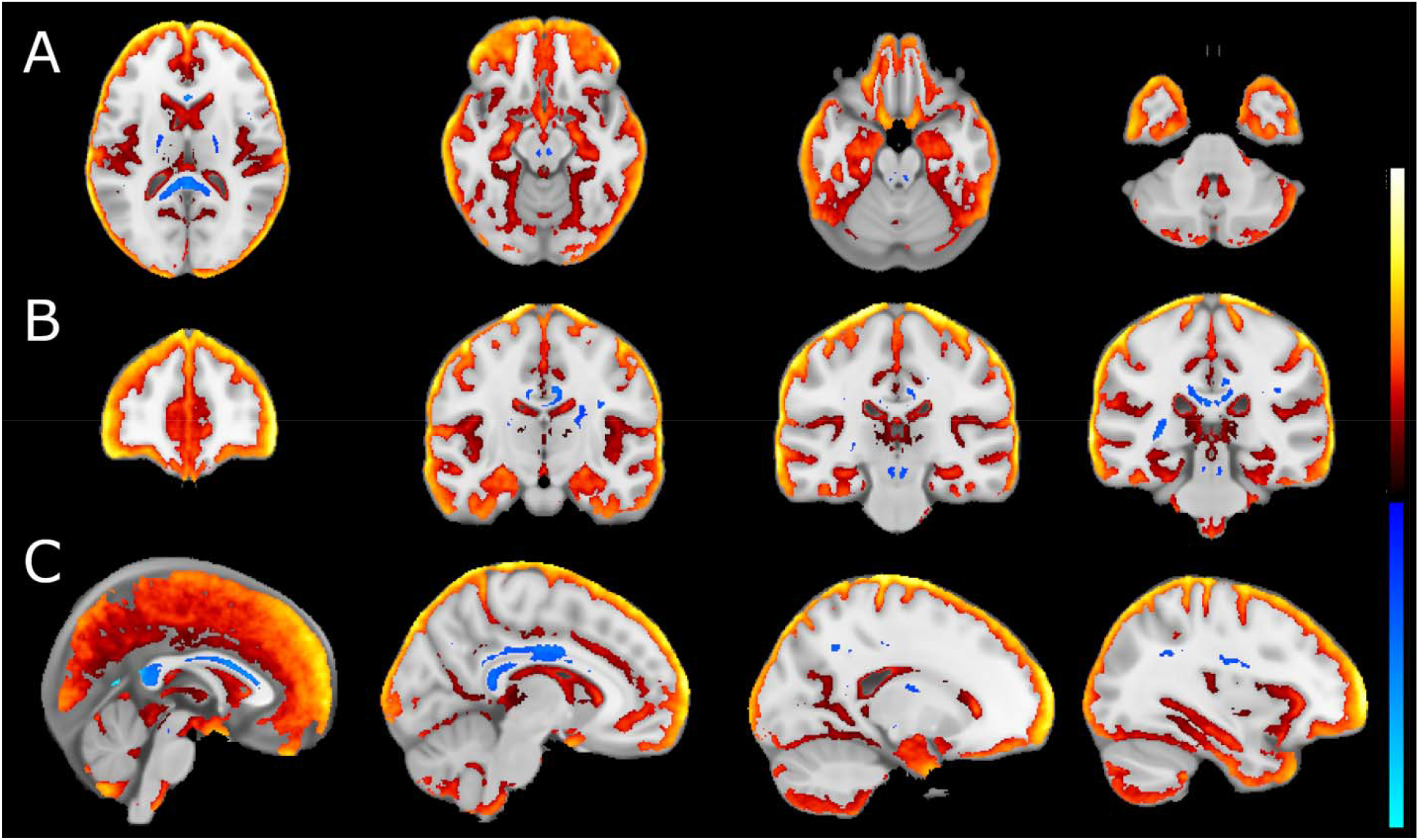
Mean diffusivity alterations in EP. (A) Axial, (B) coronal, and (C) sagittal sections of FA alterations between EP and controls. Differences in MD are estimated from the MD maps registered to MNI space and following a t-test. Hot colors represent reductions in MD, while cool colors refer to increases between healthy controls and EP cohorts larger than t = 6 and q < 0.05.

**Figure 4.**
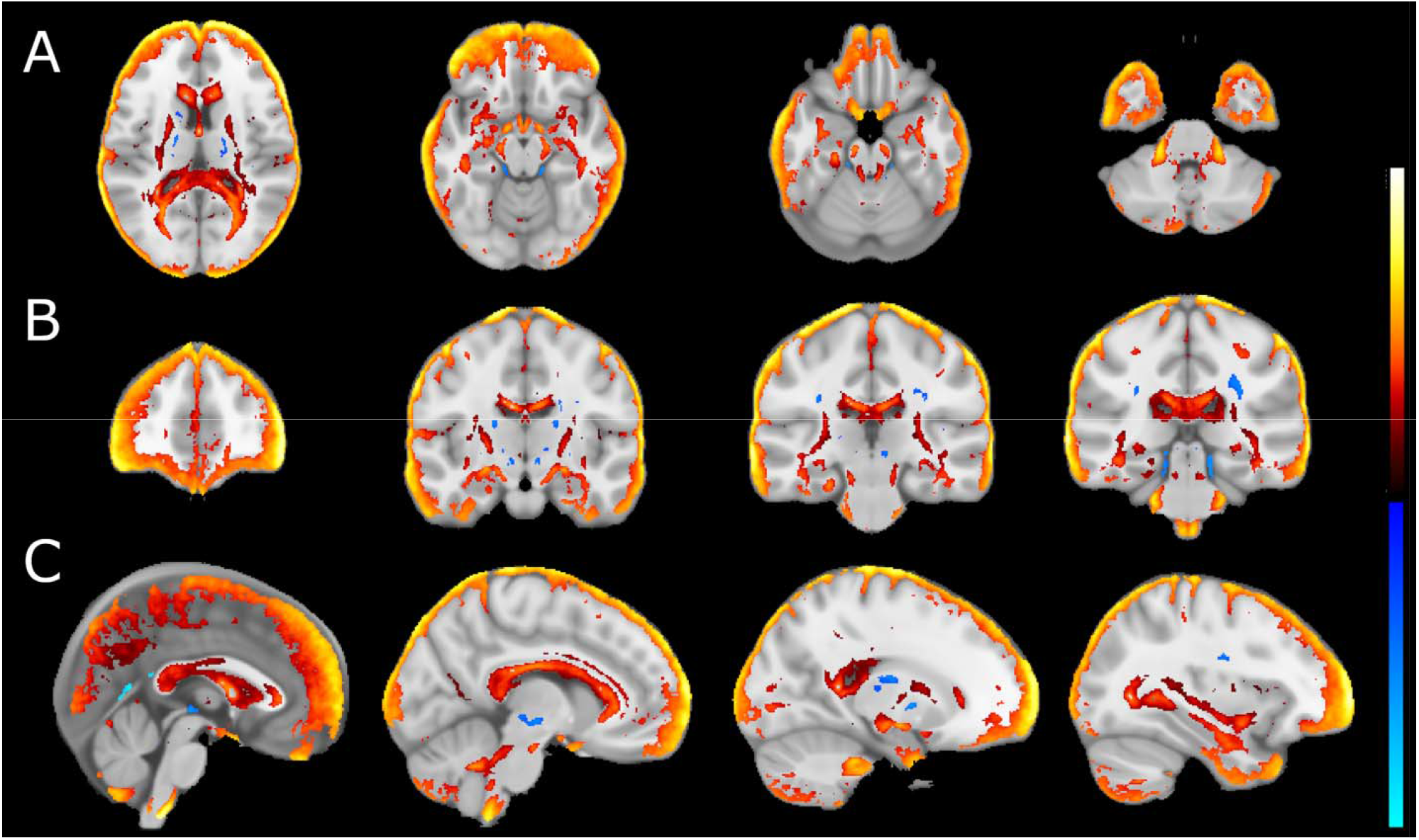
Axial diffusivity differences in EP. (A) Axial, (B) coronal, and (C) sagittal sections of AD alterations between EP and controls. Differences in AD are estimated from the AD maps registered to MNI space and following a t-test. Hot colors represent reductions in AD, while cool colors refer to increases between healthy controls and EP cohorts larger than t = 6 and q < 0.05.

In addition to the DTI indices, we carried out TBSS to extract more granular information about major white matter tracts. TBSS projects tract data onto an artificially constructed “skeleton” (represented by the green in **Fig 5)** of all white matter tracts. TBSS then performs a statistical analysis of the FA (**Supp Fig 7**) maps over this “skeleton.” TBSS results revealed significantly higher FA values in EP than controls in multiple tracks, with the highest effect sizes in the midbrain, cingulum, and corpus callosum. TBSS highlighted significant FA reductions in white matter tracts within the midbrain in EP vs. controls.

**Figure 5.**
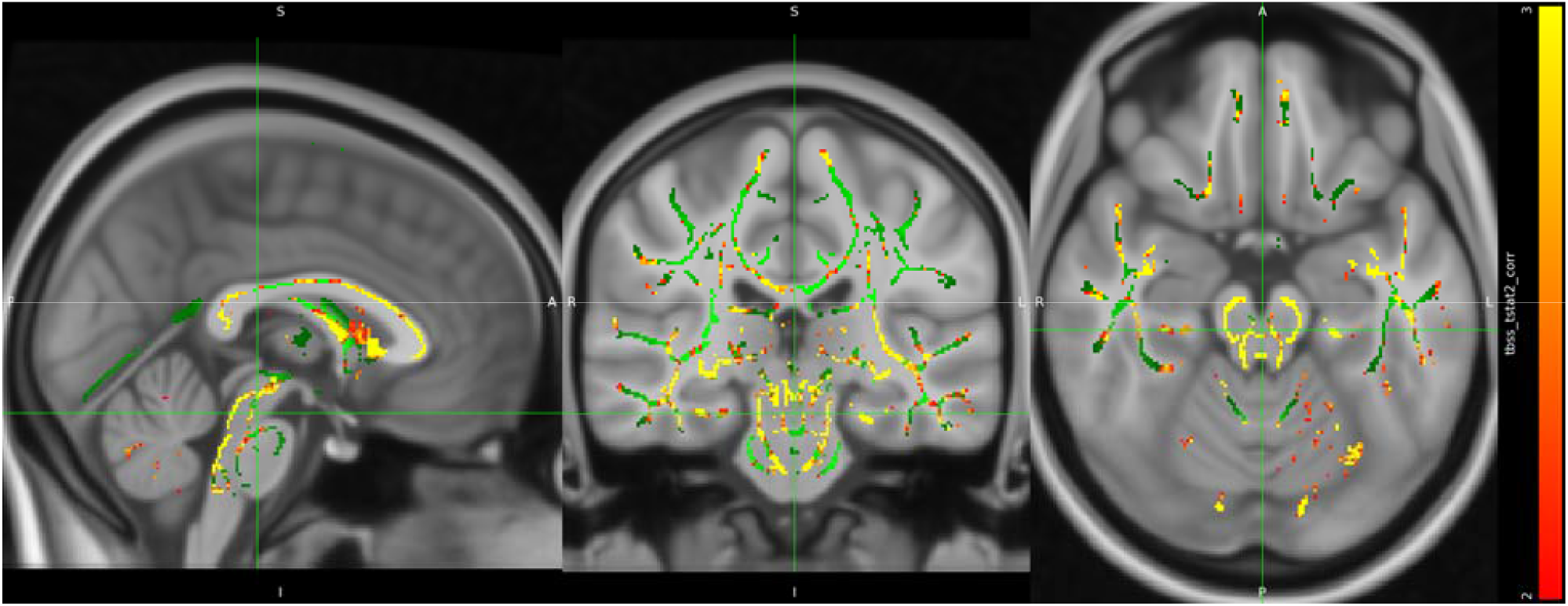
White matter alterations in EP. TBSS maps report differences from the corrected t-test following TBSS pipelines. Heat maps represent relative differences between healthy controls and EP cohorts larger than t = 2 and q < 0.05.

As the next step, a midbrain ROI was generated from FreeSurfer v.6.0. Destrieux atlas (https://mindboggle.info/data.html, ^37^. A seed-based analysis was done using the midbrain ROI on the JD, midbrain volumes, FA, MD, and AD maps to focus specifically on the midbrain macro- and micro-structure differences between EP and controls. The results showed no significant differences in the measures of JD (t = 0.96, p = 0.33) or native midbrain volumes (t = -1.61, p = 0.11). However, there were significant between-group differences in FA (t = -11.58, p = 2.2e-16), AD (t = 2.33, p = 0.02), and MD (t = 5.23, p = 1.06e-6,) in the midbrain (**Fig 6**).

**Figure 6.**
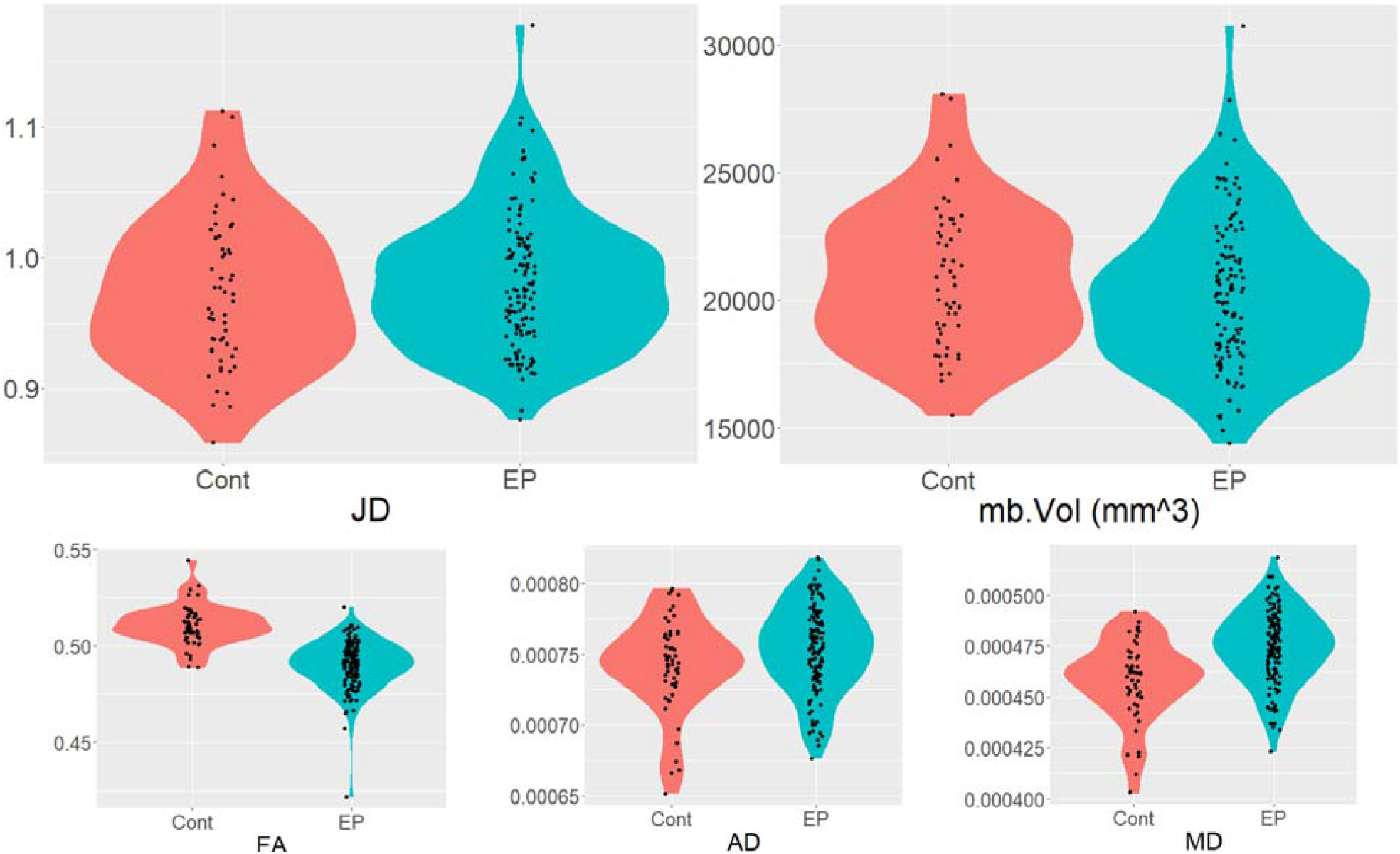
The grouped subfigures are violin plots of the distributions of midbrain microstructural alterations in EP vs. healthy controls in ROI means of JD, midbrain ROI volumes, and ROI means of FA, AD, and MD maps. And their Welch two-group t-test results, respectively.

The symptom metrics (i.e., PANSS and CAINS) displayed significant correlations between themselves, except for CAINS and the PANSS General Psychopathology subscale (r^2^ = 0.18, p = 0.056, *trend*). Among the MRI outcomes (i.e., ROI means of JD, midbrain volumes, and the mean of DTI indices), the midbrain volume negatively correlated with MD (r^2^ = -0.30, p = 0.001), AD (r^2^ = -0.24, p = 0.012) and FA (r^2^ = -0.238, p = 0.012). The midbrain AD correlated with MD (r^2^ = 0.953, p < 0.001), but FA did not correlate with either (**Fig 7, Supplementary Table 2**). The JD did not correlate with any other MRI metrics.

**Figure 7.**
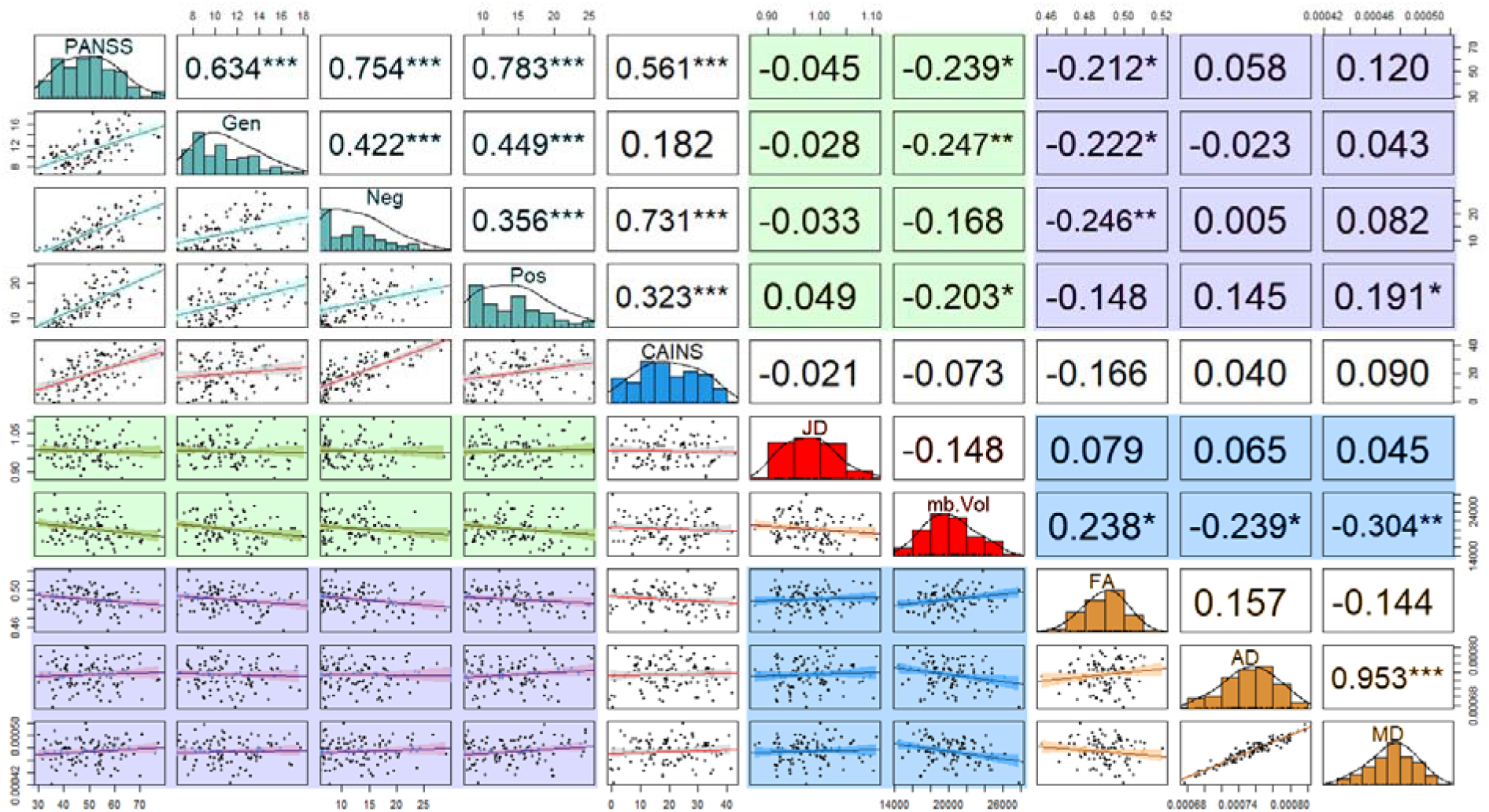
Relationships between the PANSS and CAINS symptom scores and Jacobian Determinant, midbrain volumes, and DTI indices of midbrain microstructure. The diagonal shows the distribution of each score (PANSS, CAINS, volumetrics, or DTI). Highlighted portions: Turquoise, PANSS scores; red, T1W volumetrics; orange, DTI measures; green, volumetrics vs. PANSS; purple, DTI vs PANSS; blue, volumetrics vs DTI. The bottom quadrant shows the scatter plots with fitted correlation, and the top quadrant shows r^2^ of the correlation (* = p < 0.05, ** = p < 0.01, and *** = p < 0.005).

The correlation analyses of symptom scores (i.e., PANSS, CAINS) and MRI-derived measures (i.e., volumetrics and DTI) showed significant correlations between midbrain volume and symptom severity. The midbrain volume displayed significant negative correlations with the PANSS total score (r^2^ = -0.239, p = 0.011), Positive Symptoms subscale score (r^2^ = -0.203, p = 0.032) and General Psychopathology subscale score (r^2^ = -0.247, p = 0.009); and the trend-level negative correlation with the Negative symptoms’ subscale score (r^2^ = -0.168, p = 0.077). The CAINS did not correlate with any of the MRI-derived measures. The midbrain FA showed significant negative correlations with the PANSS total score (r^2^ = -0.212, p = 0.026), General Psychopathology score (r^2^ = -0.222, p = 0.019), and Negative symptoms score (r^2^ = -0.246, p = 0.009); in addition, there was a trend for the CAINS score (r^2^ = -0.166, p = 0.081). The MD showed a significant positive correlation with the PANSS Positive symptoms score (r^2^ = 0.191, p = 0.045). The AD did not correlate with any symptom metric.

## Discussion

This study identified specific alterations in the whole brain volumetric and white matter microstructure outcomes in the publicly available HCP-EP sample. Widespread cortical and white matter alterations were identified, including in frontal, temporal, and cingulate cortical regions, corticospinal and corpus callosum white matter tracts, and midbrain. DBM shows significant differences around the midbrain and anterior hippocampus. The DTI analysis displays extensive FA, AD, and MD alterations, confirmed by a seed-based analysis of DTI metrics. The midbrain volume in EP subjects significantly correlated with several behavior indices and all DTI indices. The FA was the DTI index with the most significant correlations to behavior (i.e., PANSS overall score, general psychopathology, positive symptoms). TBSS revealed deep WM and midbrain tract alterations from EP subjects. TBSS shows alteration in the corpus callosum, cingulum, cerebral peduncles, and midbrain white matter tracts. The correlation analyses determined that FA midbrain values are associated with the severity of positive, negative, and general psychopathology PANSS-based symptoms, suggesting that the white matter microstructure alterations may be a clinically applicable biomarker of EP.

Brain structure volumetric alterations have been consistently reported in EP subjects. EP reductions left the frontotemporal region and left postcentral gyrus ^38^, hippocampus ^26,39–41^, the temporal, frontal ^42^, precentral, and insula cortex ^43^. EP subjects show lower GM volume in the left median cingulate cortex, cerebellum, thalamus, and left inferior parietal gyrus and clusters of reduced white matter ^44^. The thalamus displays volume and microstructural (assessed with DTI) alteration in EP subjects ^45,46^. GM volume reduction in EP is associated with working memory disruptions. ^47^. In a first-episode psychosis cohort, it was found that cortical thickness, surface area, and subcortical volumes display significant variability, and the researchers propose further examination to determine trajectories. ^48^. EP also displays volume reductions in the frontal area, temporal gyrus, precuneus, uncus, amygdala, and left cuneus ^49^. The EP volume alterations match those reported in schizophrenia subjects. Our results highlight medial temporal lobe alterations in the hippocampus, striatum, insula, and amygdala consistent with prior DBM findings in first-episode schizophrenia individuals ^50^.

Prior EP studies suggest that alterations in regional brain activity, gray and white matter structure, and connectivity precede and predict the psychosis onset.^12,51–53^ These early brain-based disease markers are associated with cognitive and symptom manifestations in clinical high-risk/EP populations ^54^. Young subjects with subclinical psychotic experiences (PEs) show lower mean and radial diffusivity in the arcuate fasciculus (AF) ^55^. Lower FA in regions proximal to the superior longitudinal fasciculus (SLF) and corticospinal tract bilaterally and in the left inferior frontal-occipital fasciculus (IFOF) and inferior longitudinal fasciculus (ILF) are related to higher PEs at baseline ^56^. The cerebellum shows differences in FA and radial diffusivity between clinical-high-risk (CHR) individuals and healthy controls ^57^. Psychosis risk has been linked to reductions in FA along white matter tracts connecting the limbic striatum with the limbic cortical network, the anterior right external capsule segment connections to the right prefrontal cortex ^58^. Ultra-high-risk psychosis subjects show increased radial diffusivity in the left anterior thalamic radiation and reduced bilateral thickness across the frontal, temporal, and parietal cortices ^59^. EP individuals exhibit lower generalized FA in the fornix and smaller hippocampal volumes^60^ as well as reduced mean kurtosis values in the thalamic regions and increased thalamus-orbitofrontal cortex and thalamus-lateral temporal cortex connectivity.^61^ The mean kurtosis in the thalamus and the orbitofrontal cortex correlated with spatial working memory accuracy in EP subjects; in contrast, no such significant correlations were observed in healthy subjects.^61^ There is more significant right-to-left asymmetry of the SLF in clinical high-risk for developing psychosis females compared to control females. Still, there is no hemispheric difference between clinical high-risk for developing psychosis vs. control males ^62^. Moreover, the laterality index of SLF-III for clinical high-risk developing psychosis females correlated negatively with poorer working memory functioning ^62^. These findings indicate network-level white matter microstructure alterations as a potential biomarker for CHR/EP.

DA’s role in psychotic disorders has not clearly been determined besides the fact that most approved treatments focus on blocking D2 action; however, these treatments are not always efficacious ^63^. This has set forth the idea that brain networks may modulate and be an essential target to treat psychotic disorders, particularly early in the disease. In our multimodal analyses, morphological, microstructural, and tract-based alterations in the midbrain emerged as a critical hub in EP. Prior studies point to reduced FA and higher MD in the SLF and IFOF in EP, and these are associated with hallucination severity ^64^. A network modeling highlights two trajectories of psychotic disorders, highlighting regional starting points. One trajectory starts in Broca’s area and the insula, then it propagates into frontal and medial temporal regions, then sensorimotor cortices, then in the occipital, parietal, and temporal cortices, and finally in the cerebellum and subcortical areas ^14^. The second trajectory initiates in the hippocampus and amygdala, then the parahippocampus, thalamus, and nucleus accumbens, then the caudate and insula, then the putamen, cingulate, frontal, and temporal lobes, and finally, the several cortical areas ^14^. EP subjects also display alterations in quantitative anisotropy in the corpus callosum, cerebellum, and peduncles ^65^. In addition, these alterations correlate with PANSS positive and negative symptoms ^65^. Processing speed in EP is associated with the structural connectivity of superior frontal gyri, precuneus, somatomotor, temporal-mesial cortices, thalamus, superior-parietal cortex, caudate, pallidum, and lateral-occipital cortex, and clinical subtypes of EP are characterized by distinct brain-connectivity profiles ^66^. In addition, the midbrain (i.e., its striatal system) is the primary source of dopamine synthesis and the main target of antipsychotic medication effects. The baseline connectivity of anteromedial hippocampal FC with the insular-opercular cortex, superior frontal gyrus, precentral gyrus, and postcentral gyrus predict treatment response to antipsychotics ^2^

### Study limitations and future directions

Our study presents evidence of macro- and microstructural brain alterations in the early-course phase of psychotic disorders. Captured with an assay of multimodal MRI approaches in EP’s frontal, temporal, and ^67^midbrain regions, leveraging a high-quality, large, publicly available dataset (HCP-EP). Several limitations of the study should be noted. First, the cross-sectional nature of the HCP-EP sample limits our ability to examine the structural brain alterations along the specific epochs/trajectories of evolving psychosis, e.g., “early” CHR vs. “late” CHR vs. EP. One of the strengths of our analyses is its multimodal approach, which incorporates DBM, seed-based volume analysis, voxel-based DTI analysis, and seed-based analysis of DTI metrics. However, other diffusion approaches, such as NODDI^68^ may provide more specific markers of the microstructural basis of the alterations in DTI metrics. This study focused on brain structure alterations in EP; hence, the potential alterations in functional networks have not been examined. Our future work will expand these findings to 1) determine the topological alterations in brain structure using tractography and volume analysis of white matter tracts associated with psychotic disorders, 2) sex differences in EP, and 3) functional connectivity analysis of resting state networks.

## Data availability

Data can be obtained through an NDA with NIMH. Instruction and relevant information can be found https://www.humanconnectome.org/study/human-connectome-project-for-early-psychosis/document/hcp-ep

## Declarations

Data used in the preparation of this manuscript was obtained from the National Institute of Mental Health (NIMH) Data Archive (NDA). NDA is a collaborative informatics system created by the National Institutes of Health to provide a national resource to support and accelerate research in mental health. Dataset identifier(s): NDA Collection #2914. We obtained a Data Use Certification to use the HCP-EP data set (OMB Control Number: 0925-0667) and comply with all safety and regulations for using the data. This manuscript reflects the views of the authors and may not reflect the opinions or views of the NIH or of the Submitters submitting original data to NDA.

## Acknowledgment

LMCP was supported by NIH/NIDA award K25 during the preparation of this manuscript. EII was supported by NIH/NIMH award R01MH127317 during the preparation of the manuscript. The NIH/NIMH had no further role in study design, data analyses, and interpretation of findings in the writing of the manuscript or in the decision to submit the manuscript for publication.

## References

1. Shinn, A. K. et al. McLean OnTrack: a transdiagnostic program for early intervention in first-episode psychosis. Early Interv Psychiatry 11, 83–90 (2017).

2. Blessing, E. M. et al. Anterior Hippocampal-Cortical Functional Connectivity Distinguishes Antipsychotic Naïve First-Episode Psychosis Patients From Controls and May Predict Response to Second-Generation Antipsychotic Treatment. Schizophr Bull 46, 680–689 (2020).

3. Kane, J. M. et al. Comprehensive versus usual community care for first-episode psychosis: 2-Year outcomes from the NIMH RAISE early treatment program. American Journal of Psychiatry 173, 362–372 (2016).

4. Xiao, Y. et al. White Matter Abnormalities in Never-Treated Patients With Long-Term Schizophrenia. Am J Psychiatry 175, 1129–1136 (2018).

5. Hardy, K. V, Association, A. P., Ballon, J. S., Noordsy, D. L. & Adelsheim, S. Intervening Early in Psychosis[]: A Team Approach. First edit, (2019).

6. Blackman, G. et al. Prevalence of Neuroradiological Abnormalities in First-Episode Psychosis: A Systematic Review and Meta-analysis. JAMA Psychiatry 80, 1047–1054 (2023).

7. Ji, J. L. et al. Mapping brain-behavior space relationships along the psychosis spectrum. Elife 10, (2021).

8. O’Neill, A., Mechelli, A. & Bhattacharyya, S. Dysconnectivity of Large-Scale Functional Networks in Early Psychosis: A Meta-analysis. Schizophr Bull 45, 579 (2019).

9. Braun, U. et al. Dynamic brain network reconfiguration as a potential schizophrenia genetic risk mechanism modulated by NMDA receptor function. Proc Natl Acad Sci U S A 113, 12568–12573 (2016).

10. Rubinov, M. & Bullmore, E. Schizophrenia and abnormal brain network hubs. Dialogues Clin Neurosci 15, 339 (2013).

11. Chopra, S. et al. Network-Based Spreading of Gray Matter Changes Across Different Stages of Psychosis. JAMA Psychiatry 80, 1246–1257 (2023).

12. Schobel, S. A. et al. Imaging Patients with Psychosis and a Mouse Model Establishes a Spreading Pattern of Hippocampal Dysfunction and Implicates Glutamate as a Driver. Neuron 78, 81–93 (2013).

13. Cannon, T. D. et al. Progressive reduction in cortical thickness as psychosis develops: a multisite longitudinal neuroimaging study of youth at elevated clinical risk. Biol Psychiatry 77, 147–157 (2015).

14. Jiang, Y. et al. Neuroimaging biomarkers define neurophysiological subtypes with distinct trajectories in schizophrenia. Nature Mental Health 2023 1:3 1, 186–199 (2023).

15. Allen, P. et al. Resting hyperperfusion of the hippocampus, midbrain, and basal ganglia in people at high risk for psychosis. American Journal of Psychiatry 173, 392–399 (2016).

16. Romaniuk, L. et al. Midbrain Activation During Pavlovian Conditioning and Delusional Symptoms in Schizophrenia. Arch Gen Psychiatry 67, 1246–1254 (2010).

17. Gangadin, S. S., Cahn, W., Scheewe, T. W., Hulshoff Pol, H. E. & Bossong, M. G. Reduced resting state functional connectivity in the hippocampus-midbrain-striatum network of schizophrenia patients. J Psychiatr Res 138, 83–88 (2021).

18. Howes, O. D. et al. Midbrain dopamine function in schizophrenia and depression: a post-mortem and positron emission tomographic imaging study. Brain 136, 3242–3251 (2013).

19. Modinos, G., Allen, P., Grace, A. A. & McGuire, P. Translating the MAM Model of Psychosis to Humans. Trends Neurosci 38, 129 (2015).

20. Lisman, J. E. et al. Circuit-based framework for understanding neurotransmitter and risk gene interactions in schizophrenia. Trends Neurosci 31, 234–242 (2008).

21. Lodge, D. J. & Grace, A. A. Hippocampal dysregulation of dopamine system function and the pathophysiology of schizophrenia. Trends Pharmacol Sci 32, 507–513 (2011).

22. Mahdiar, M. et al. Raphe Nuclei Echogenicity and Diameter of Third Ventricle in Schizophrenia Measured by Transcranial Sonography. Basic Clin Neurosci 14, 463–470 (2023).

23. Purves-Tyson, T. D. et al. Increased levels of midbrain immune-related transcripts in schizophrenia and in murine offspring after maternal immune activation. Molecular Psychiatry 2019 26:3 26, 849–863 (2019).

24. Bielawski, M. & Bondurant, H. Psychosis following a stroke to the cerebellum and midbrain: a case report. Cerebellum Ataxias 2, (2015).

25. Gregory, D. F. et al. Increased functional coupling between VTA and hippocampus during rest in first-episode psychosis. eNeuro 8, 1–8 (2021).

26. McHugo, M. et al. Anterior hippocampal dysfunction in early psychosis: A 2-year follow-up study. Psychol Med (2021) doi:10.1017/S0033291721001318.

27. American Psychiatric Association. Diagnostic and Statistical Manual of Mental Disorders. Diagnostic and Statistical Manual of Mental Disorders (2022) doi:10.1176/APPI.BOOKS.9780890425787.

28. Ashburner, J. & Friston, K. J. Voxel-based morphometry--the methods. Neuroimage 11, 805–821 (2000).

29. Marcus, D. S., Fotenos, A. F., Csernansky, J. G., Morris, J. C. & Buckner, R. L. Open Access Series of Imaging Studies (OASIS): Longitudinal MRI Data in Nondemented and Demented Older Adults. J Cogn Neurosci 22, 2677 (2010).

30. Antonopoulos, G. et al. A systematic comparison of VBM pipelines and their application to age prediction. Neuroimage 279, (2023).

31. Chung, M. K. et al. A unified statistical approach to deformation-based morphometry. Neuroimage 14, 595–606 (2001).

32. Smith, S. M. et al. Tract-based spatial statistics: voxelwise analysis of multi-subject diffusion data. Neuroimage 31, 1487–1505 (2006).

33. Kay, S. R., Fiszbein, A. & Opler, L. A. The positive and negative syndrome scale (PANSS) for schizophrenia. Schizophr Bull 13, 261–276 (1987).

34. Kring, A. M., Gur, R. E., Blanchard, J. J., Horan, W. P. & Reise, S. P. The Clinical Assessment Interview for Negative Symptoms (CAINS): final development and validation. Am J Psychiatry 170, 165–172 (2013).

35. Kay, S. R., Opler, L. A. & Lindenmayer, J. P. Reliability and validity of the positive and negative syndrome scale for schizophrenics. Psychiatry Res 23, 99–110 (1988).

36. Manera, A. L., Dadar, M., Collins, D. L. & Ducharme, S. Deformation based morphometry study of longitudinal MRI changes in behavioral variant frontotemporal dementia. Neuroimage Clin 24, 102079 (2019).

37. Klein, A. & Tourville, J. 101 labeled brain images and a consistent human cortical labeling protocol. Front Neurosci 6, (2012).

38. Kolenič, M. et al. Higher Body-Mass Index and Lower Gray Matter Volumes in First Episode of Psychosis. Front Psychiatry 11, (2020).

39. McHugo, M. et al. Smaller anterior hippocampal subfields in the early stage of psychosis. Transl Psychiatry 14, (2024).

40. Brunner, G. et al. Hippocampal structural alterations in early-stage psychosis: Specificity and relationship to clinical outcomes. Neuroimage Clin 35, (2022).

41. Briend, F. et al. Hippocampal glutamate and hippocampus subfield volumes in antipsychotic-naive first episode psychosis subjects and relationships to duration of untreated psychosis. Transl Psychiatry 10, (2020).

42. Yang, K. et al. A multimodal study of a first episode psychosis cohort: potential markers of antipsychotic treatment resistance. Mol Psychiatry 27, 1184–1191 (2022).

43. Garcia-Marti, G. et al. Progressive loss of cortical gray matter in first episode psychosis patients with auditory hallucinations. Schizophr Res (2023) doi:10.1016/J.SCHRES.2023.11.011.

44. Si, S. et al. Mapping gray and white matter volume abnormalities in early-onset psychosis: an ENIGMA multicenter voxel-based morphometry study. Mol Psychiatry (2024) doi:10.1038/S41380-023-02343-1.

45. Alemán-Gómez, Y. et al. Multimodal Magnetic Resonance Imaging Depicts Widespread and Subregion Specific Anomalies in the Thalamus of Early-Psychosis and Chronic Schizophrenia Patients. Schizophr Bull 49, 196–207 (2023).

46. Alemán-Gómez, Y. et al. Partial-volume modeling reveals reduced gray matter in specific thalamic nuclei early in the time course of psychosis and chronic schizophrenia. Hum Brain Mapp 41, 4041–4061 (2020).

47. Rapado-Castro, M. et al. Fronto-Parietal Gray Matter Volume Loss Is Associated with Decreased Working Memory Performance in Adolescents with a First Episode of Psychosis. J Clin Med 10, (2021).

48. Antoniades, M. et al. Personalized Estimates of Brain Structural Variability in Individuals With Early Psychosis. Schizophr Bull 47, 1029–1038 (2021).

49. Canal-Rivero, M. et al. Brain grey matter abnormalities in first episode non-affective psychosis patients with suicidal behaviours: The role of neurocognitive functioning. Prog Neuropsychopharmacol Biol Psychiatry 102, (2020).

50. Torres, U. S. et al. Patterns of regional gray matter loss at different stages of schizophrenia: A multisite, cross-sectional VBM study in first-episode and chronic illness. Neuroimage Clin 12, 1–15 (2016).

51. Lewandowski, K., Bouix, S., Ongur, D. & Shenton, M. Neuroprogression across the Early Course of Psychosis. J Psychiatr Brain Sci 5, (2020).

52. Keshavan, M. S., DeLisi, L. E. & Seidman, L. J. Early and broadly defined psychosis risk mental states. Schizophr Res 126, 1–10 (2011).

53. Cao, H. et al. Altered Brain Activation During Memory Retrieval Precedes and Predicts Conversion to Psychosis in Individuals at Clinical High Risk. Schizophr Bull 45, 924–933 (2019).

54. Jalbrzikowski, M. et al. Association of Structural Magnetic Resonance Imaging Measures With Psychosis Onset in Individuals at Clinical High Risk for Developing Psychosis: An ENIGMA Working Group Mega-analysis. JAMA Psychiatry 78, 753–766 (2021).

55. Dooley, N. et al. Psychotic experiences in childhood are associated with increased structural integrity of the left arcuate fasciculus – A population-based case-control study. Schizophr Res 215, 378–384 (2020).

56. DeRosse, P. et al. White matter abnormalities associated with subsyndromal psychotic-like symptoms predict later social competence in children and adolescents. Schizophr Bull 43, 152– 159 (2017).

57. Fitzsimmons, J. et al. Cingulum bundle abnormalities and risk for schizophrenia. Schizophr Res 215, 385–391 (2020).

58. Straub, K. T., Hua, J. P. Y., Karcher, N. R. & Kerns, J. G. Psychosis risk is associated with decreased white matter integrity in limbic network corticostriatal tracts. Psychiatry Res Neuroimaging 301, (2020).

59. Tomyshev, A. S. et al. Alterations in white matter microstructure and cortical thickness in individuals at ultra-high risk of psychosis: A multimodal tractography and surface-based morphometry study. Psychiatry Res Neuroimaging 289, 26–36 (2019).

60. Baumann, P. S. et al. Impaired fornix–hippocampus integrity is linked to peripheral glutathione peroxidase in early psychosis. Transl Psychiatry 6, e859–e859 (2016).

61. Cho, K. I. K. et al. Microstructural Changes in Higher-Order Nuclei of the Thalamus in Patients With First-Episode Psychosis. Biol Psychiatry 85, 70–78 (2019).

62. Steinmann, S. et al. Sex-Related Differences in White Matter Asymmetry and Its Implications for Verbal Working Memory in Psychosis High-Risk State. Front Psychiatry 12, (2021).

63. Howes, O. D., Bukala, B. R. & Beck, K. Schizophrenia: from neurochemistry to circuits, symptoms and treatments. Nat Rev Neurol (2023) doi:10.1038/S41582-023-00904-0.

64. Sato, Y. et al. Relationship Between White Matter Microstructure and Hallucination Severity in the Early Stages of Psychosis: A Diffusion Tensor Imaging Study. Schizophr Bull Open 2, (2021).

65. Moghaddam, H. S. et al. White matter alterations in affective and non-affective early psychosis: A diffusion MRI study. J Affect Disord 351, 615–623 (2024).

66. Griffa, A. et al. Brain connectivity alterations in early psychosis: from clinical to neuroimaging staging. Transl Psychiatry 9, (2019).

67. Keshavan, M. S. & Amirsadri, A. Early intervention in schizophrenia: Current and future perspectives. Current Psychiatry Reports vol. 9 325–328 Preprint at 10.1007/s11920-007-0040-8 (2007).

68. Zhang, H., Schneider, T., Wheeler-Kingshott, C. A. & Alexander, D. C. NODDI: practical in vivo neurite orientation dispersion and density imaging of the human brain. Neuroimage 61, 1000– 1016 (2012).

