## Supplementary figures and tables for "Macro- and Micro-Structural Alterations in the Midbrain in Early Psychosis"

**Early psychosis structural abnormalities in the midbrain correlate with positive and negative symptoms --- Supplementary Materials**

Zicong Zhou^1^, Kylie Jones^1^, Elena Ivleva^2^, and Luis Colon-Perez^1*^

^1^ Department of Pharmacology and Neuroscience, University of North Texas Health Science Center, 3500 Camp Bowie Blvd, Fort Worth, TX, 76107, United States of America

^2^ Department of Psychiatry, Southwestern Medical Center, University of Texas, Dallas, TX, 75390, United States of America

**Running title: Early psychosis correlates with positive and negative symptoms.**

* Corresponding author:

Luis Colon-Perez, PhD

University of North Texas Health Science Center

3500 Camp Bowie Boulevard

Fort Worth, TX 76107

817-735-7679

**Tables.**

|  | EP: 183 | Controls: 68 | All: 251 |
| --- | --- | --- | --- |
| Age (yrs.) | 23.06 (3.64) | 24.43 (4.27) | 23.44 (3.86) |
| Gender | M: 114; F: 70 | M: 4; F: 24 | M: 157; F: 94 |
| Handedness | R: 157; L: 17; B: 8  N/A: 1 | R: 56; L: 11; B: 1 | R: 213; L: 28; B: 9  N/A: 1 |
| Psychosis Phenotype | Affective: 57  Non-affective: 126 | NA | - |
| Lifetime Antipsychotic drug (mons.) | Yes: 133, No: 49  12.42 (15.01) | NA | Yes: 134, No: 49  9.10 (13.93) |
| Ethnicity | Asian: 12  Black: 59  Hawaiian: 1  Hispanic/Latino: 4  Indigenous: 1  Multi-racial: 15  White: 91 | Asian: 10  Black: 7  Multi-racial: 9  White: 41  N/A: 1 | Asian: 22  Black: 66  Hawaiian: 1  Hispanic/Latino: 4  Indigenous: 1  Multi-racial: 24  White: 132  N/A: 1 |
| Education (in degrees and certificates earned) | 6.53 (2.09) | 8.66 (2.30) | 7.11 (2.34) |
| IQ | 103.77 (16.20) | 114.96 (10.88) | 106.80 (15.73) |

Table 1. Demographics of the HCP-EP data set.

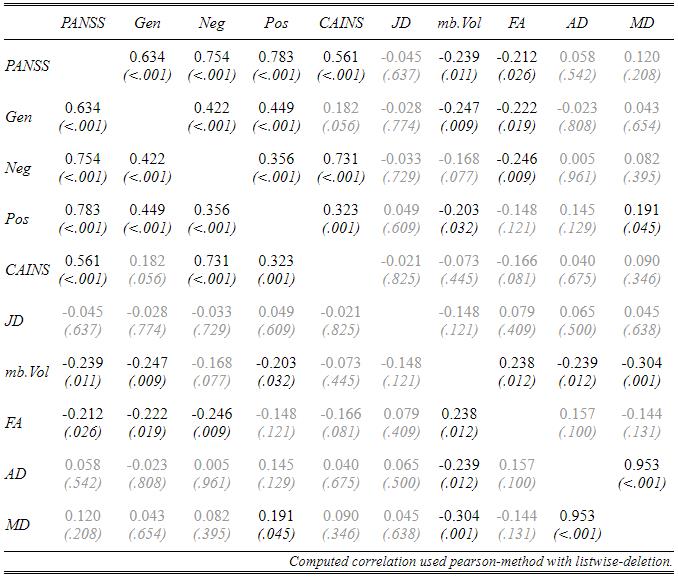

Table. 2. Correlations results of Fig 7.

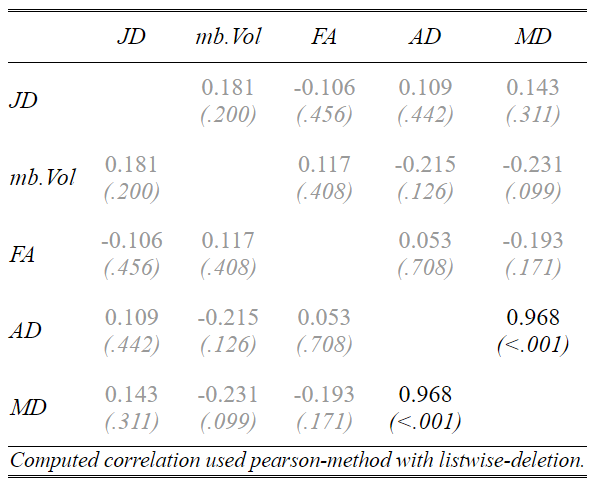

Table 3. Correlations between MRI derived metrics on the control population

Figures.

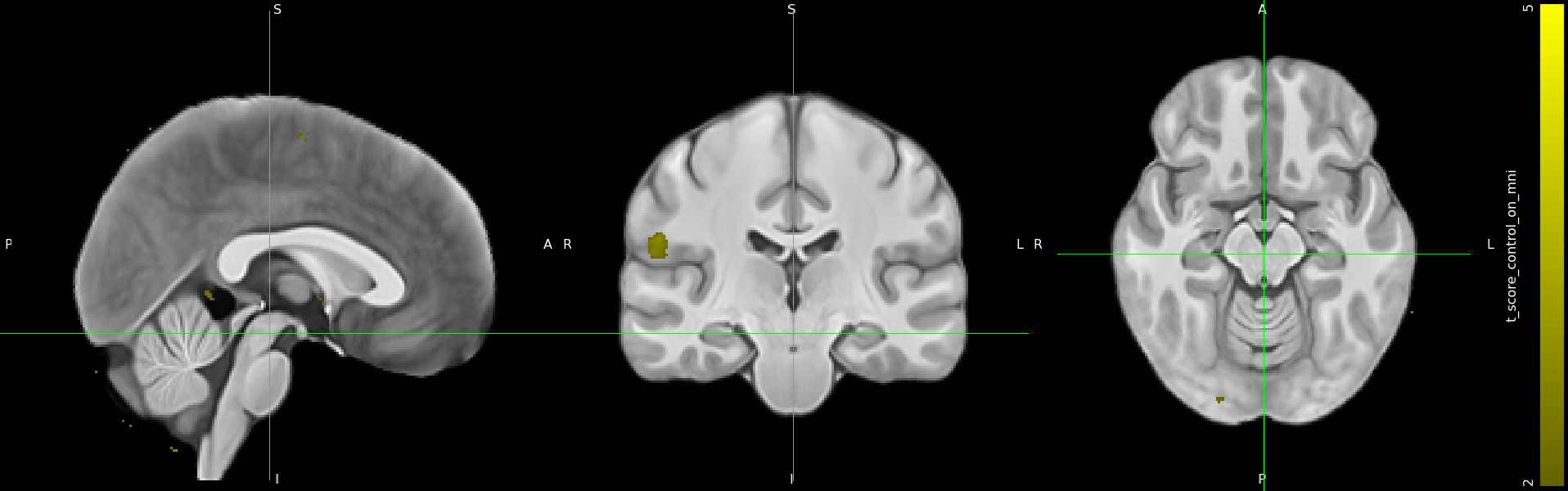

Fig 1. VBM based on the local template displayed on MNI152 space.

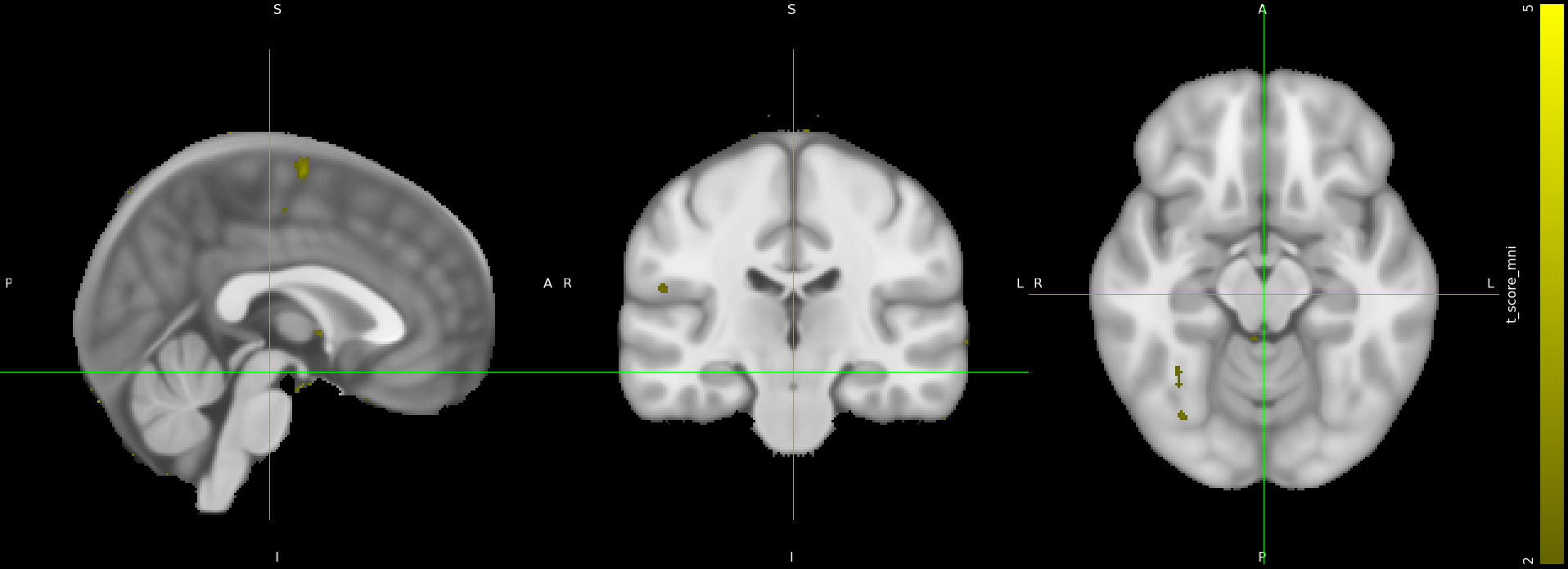

Fig 2. VBM based on MNI152

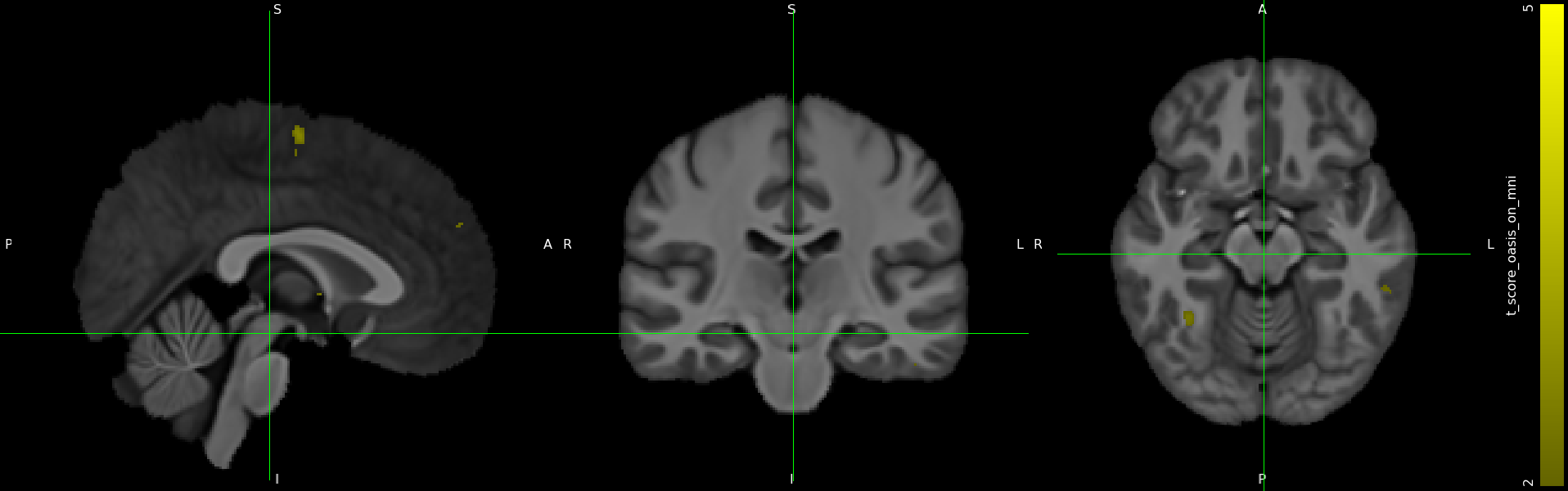

Fig 3. VBM based on OASIS template displayed on MNI152 space

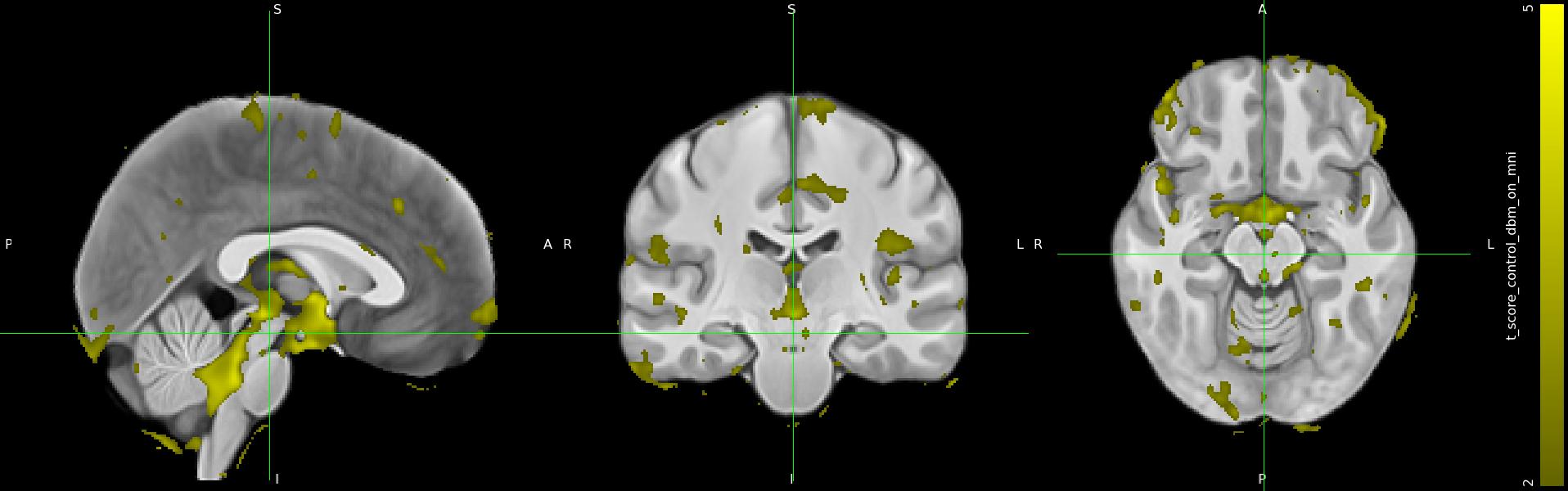

Fig 4. DBM based on the local template displayed on MNI152 space

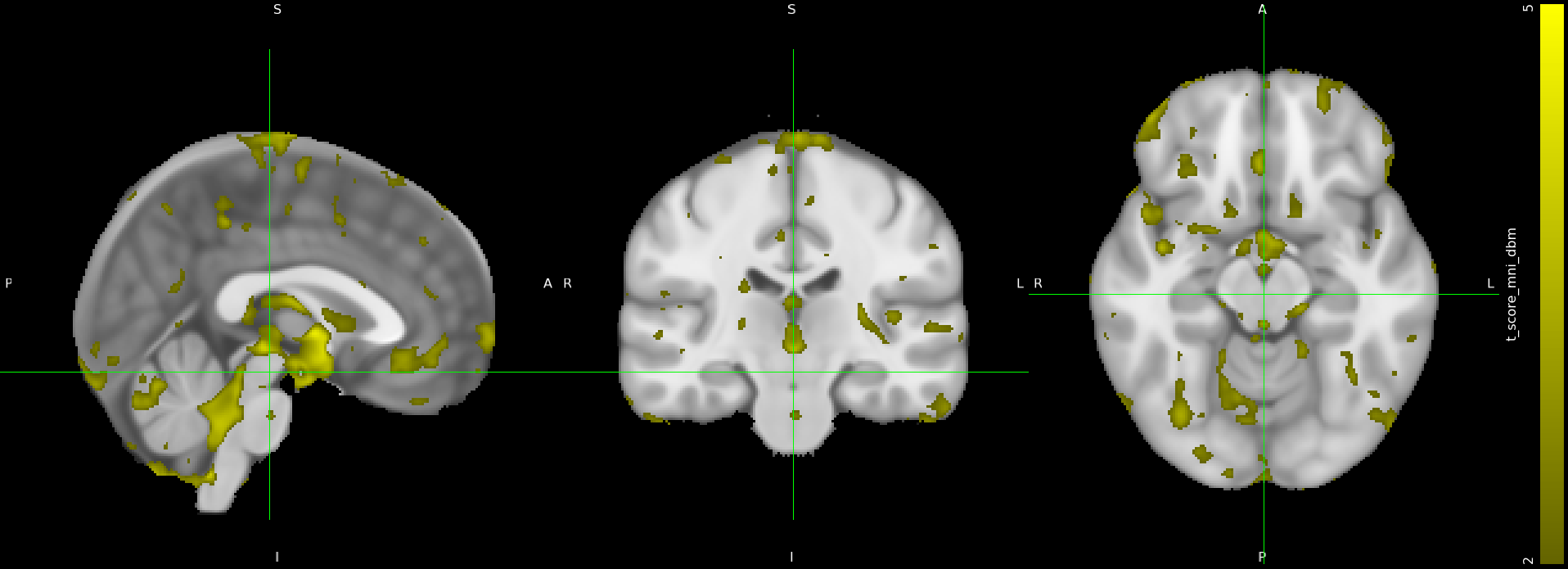

Fig 5. DBM based on MNI152 space

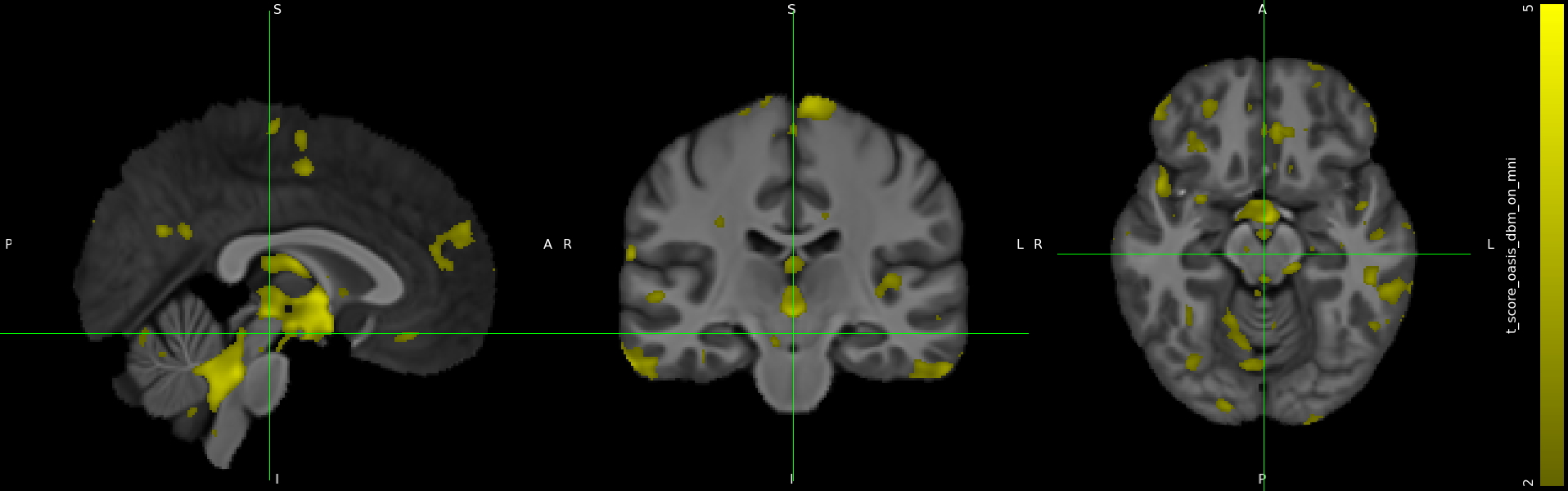

Fig 6. DBM based on OASIS template displayed on MNI152 space

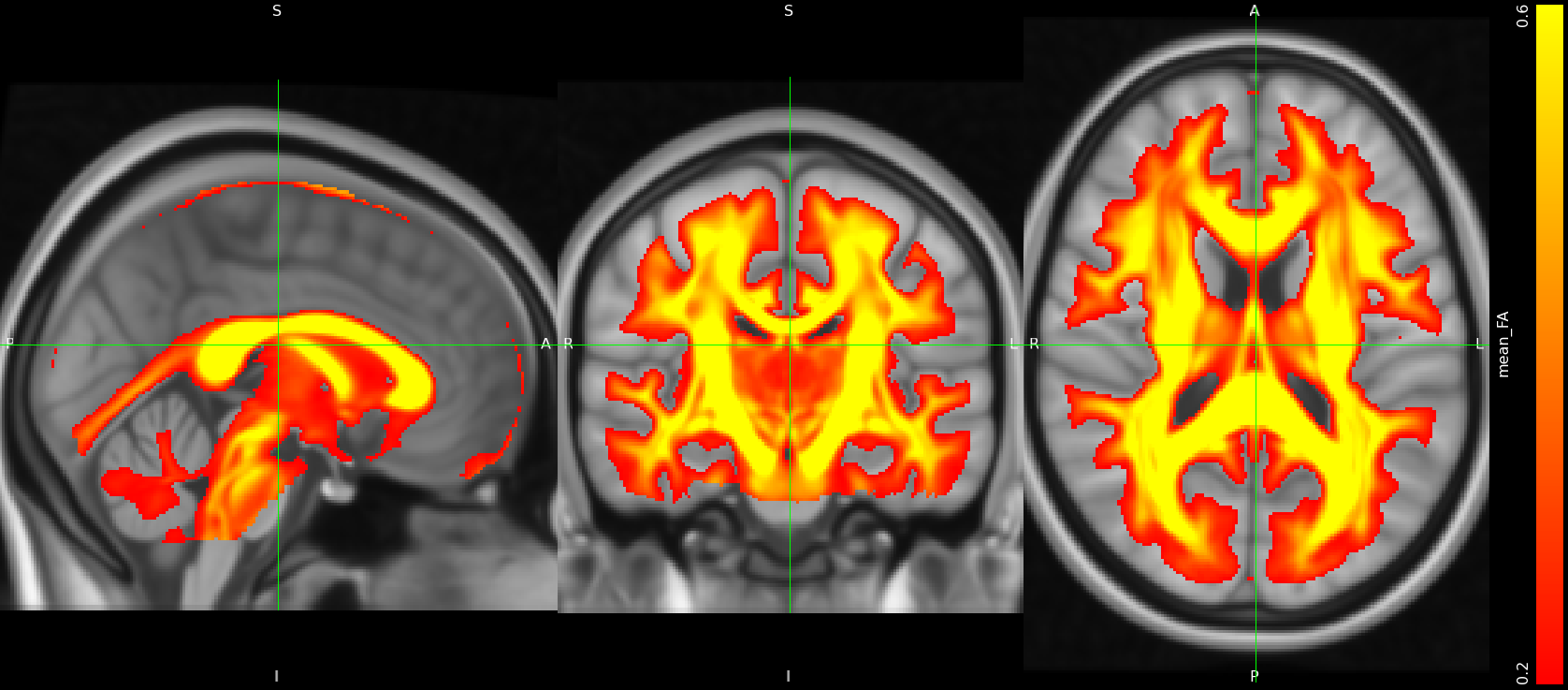

Fig 7. The mean value of all FA maps

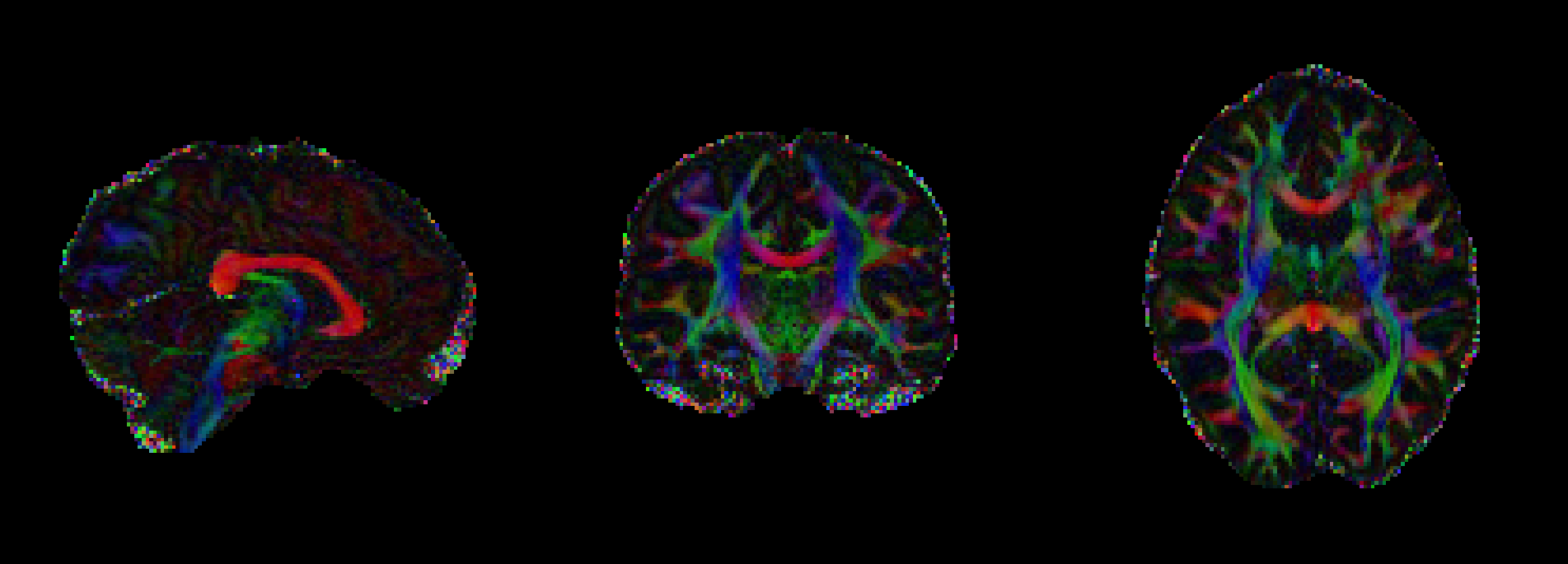

Fig 8. FA map visualization colored by its corresponding eigenvectors

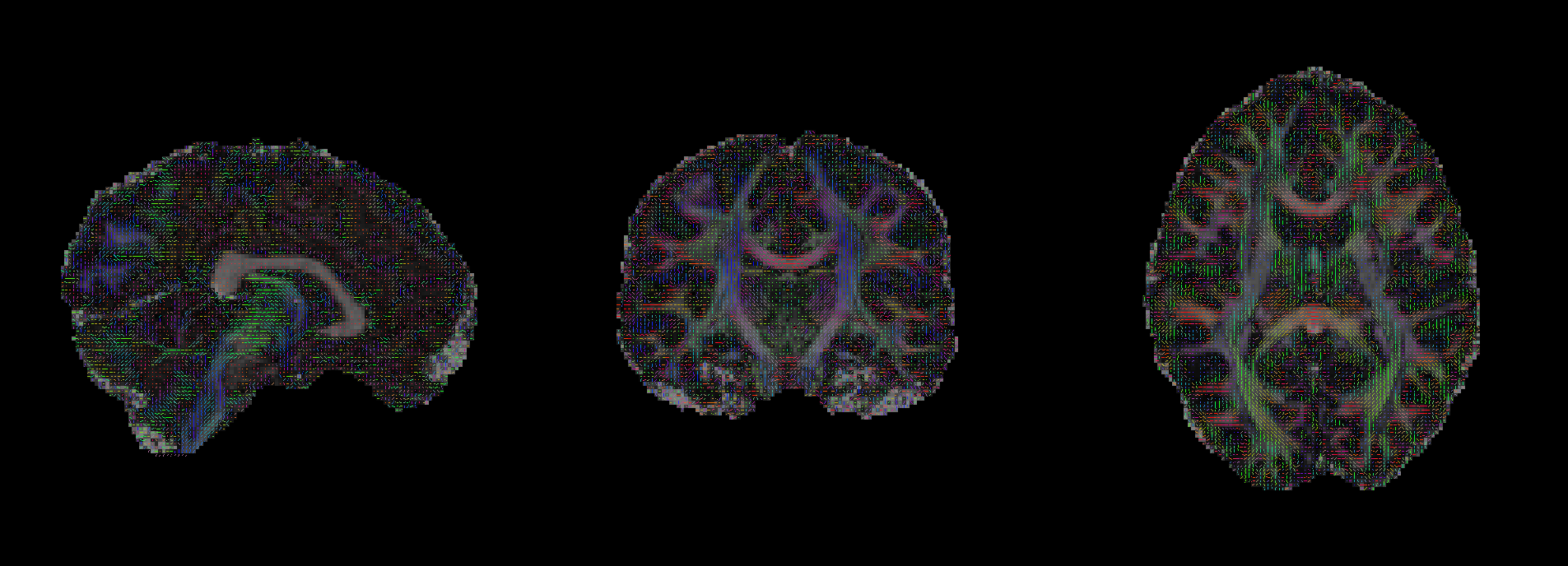

Fig 9. FA map visualization line segments by its corresponding eigenvectors

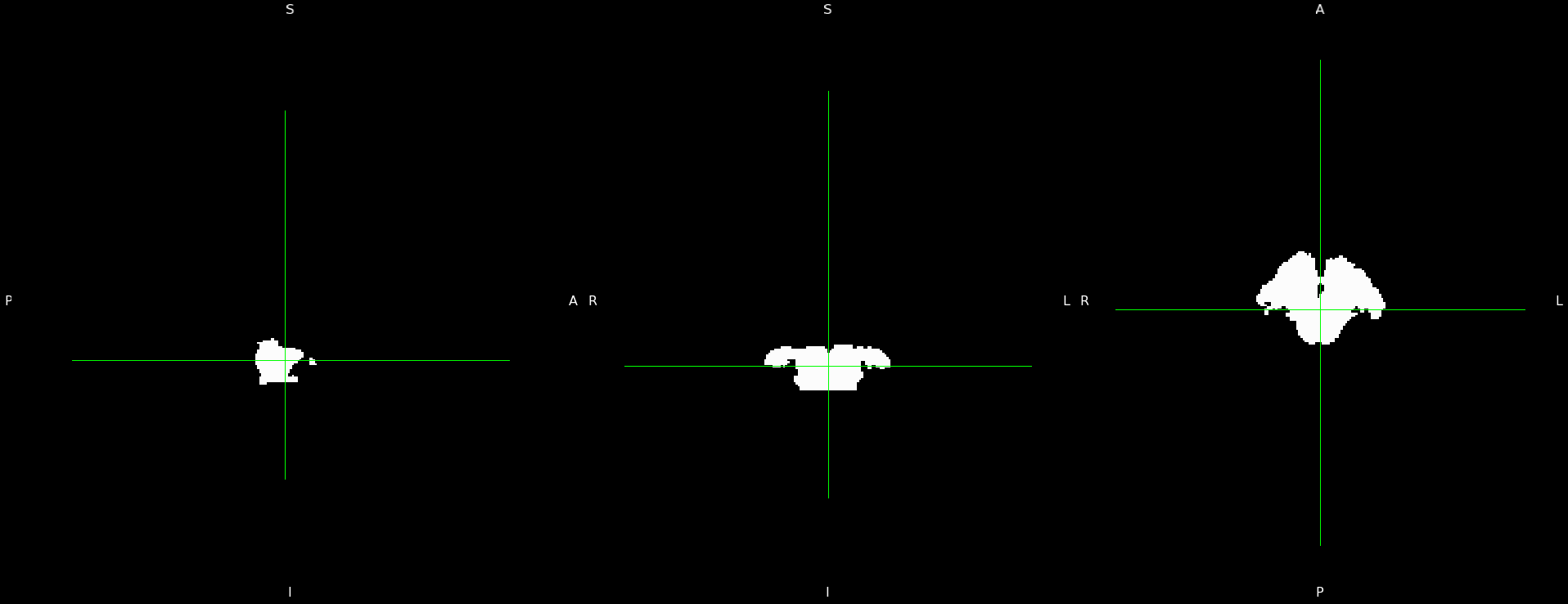

Fig 10. Midbrain Mask generated based on Freesurfer altas ROI labels

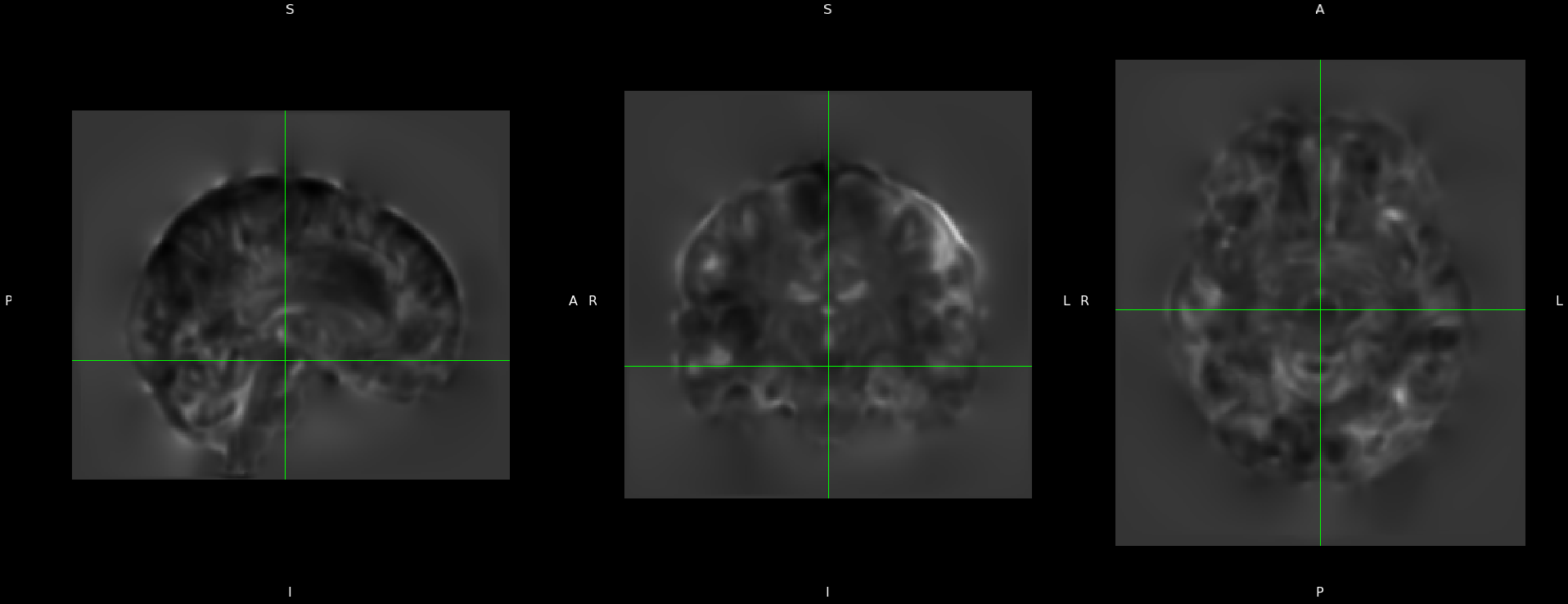

Fig 11. JD of the registration deformation from patient sample to MNI152 template

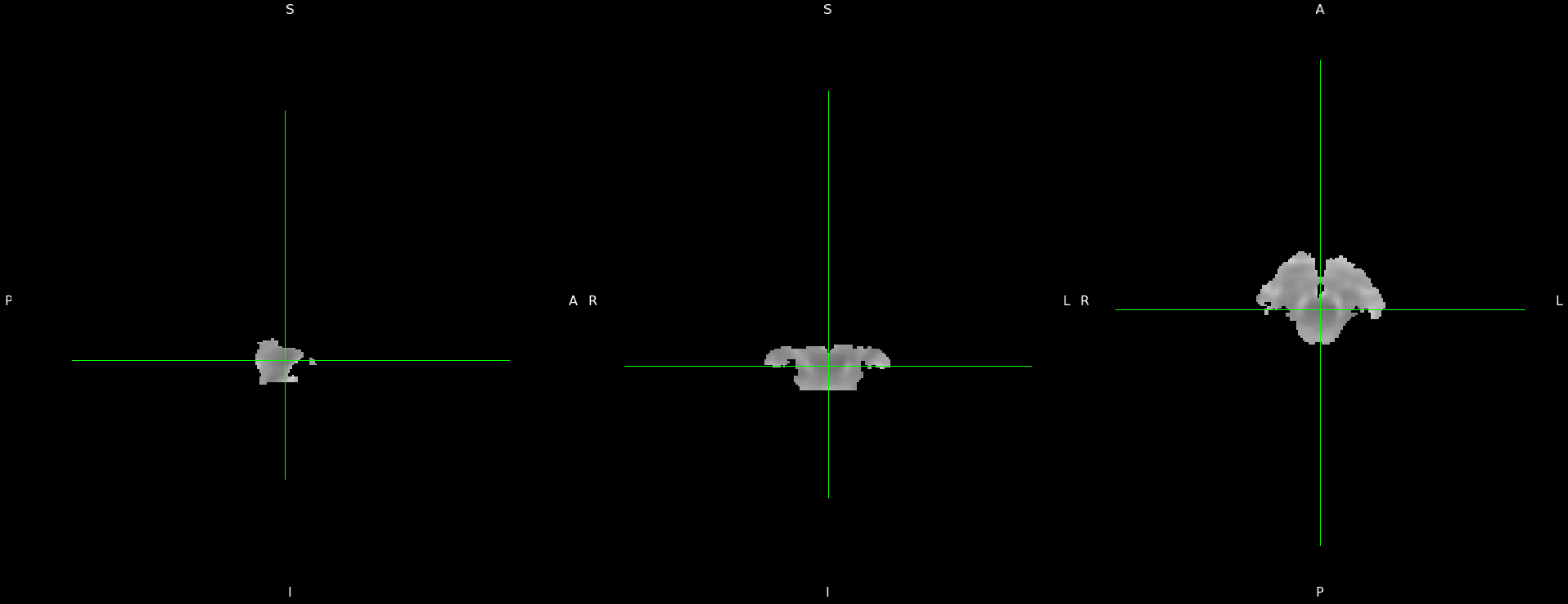

Fig 12. Midbrain segmentation of JD (brightness was scaled for visualization, values unchanged)

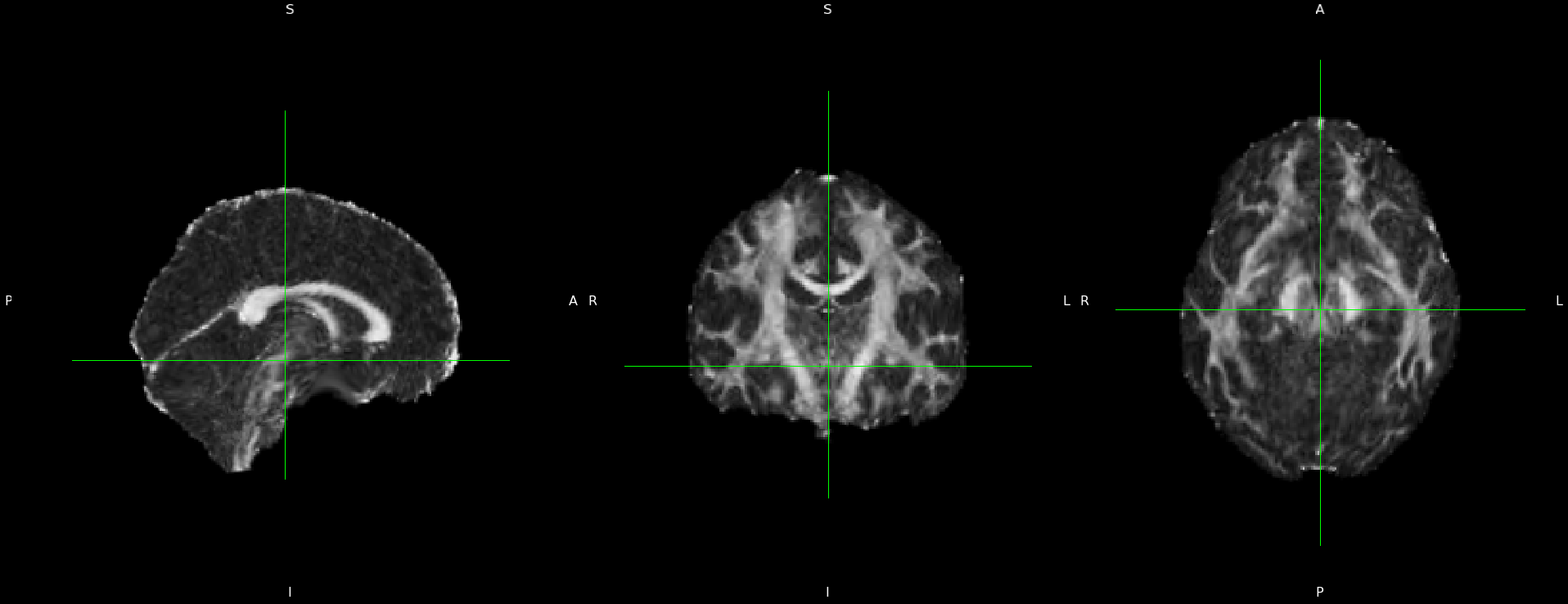

Fig 13. FA map of patient sample

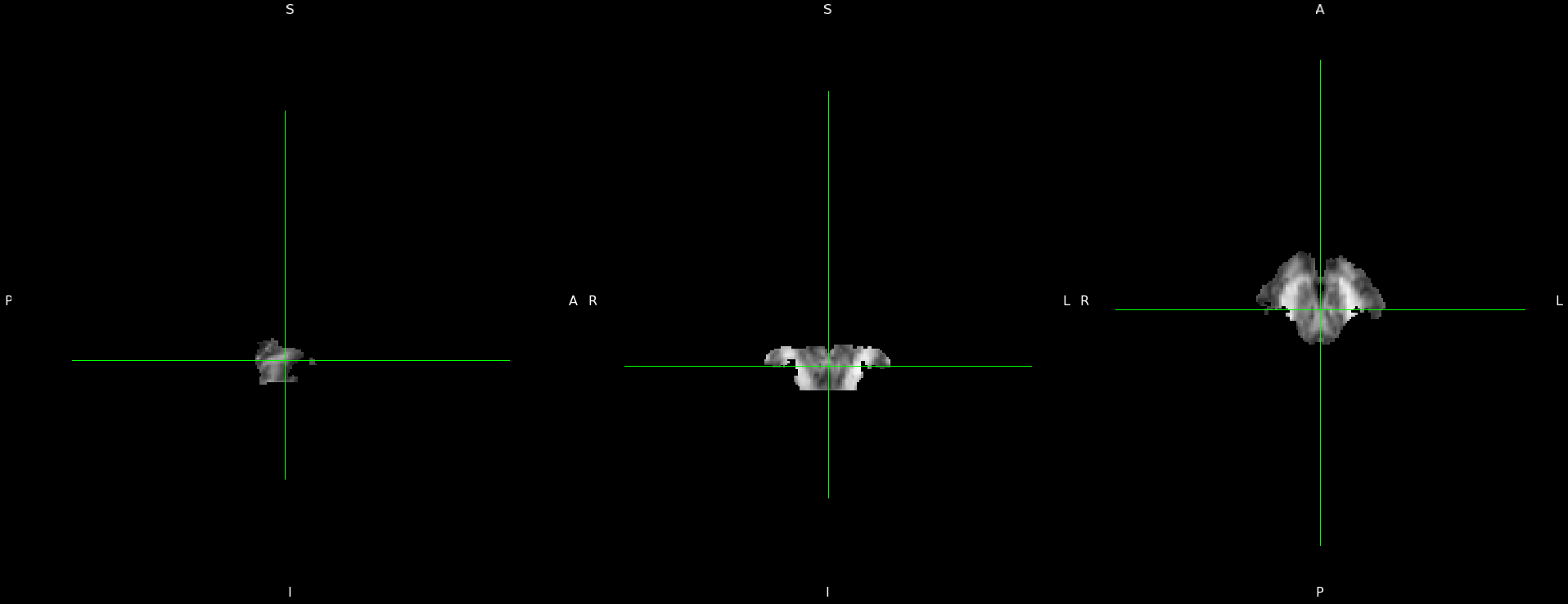

Fig 14. Midbrain segmentation of FA map

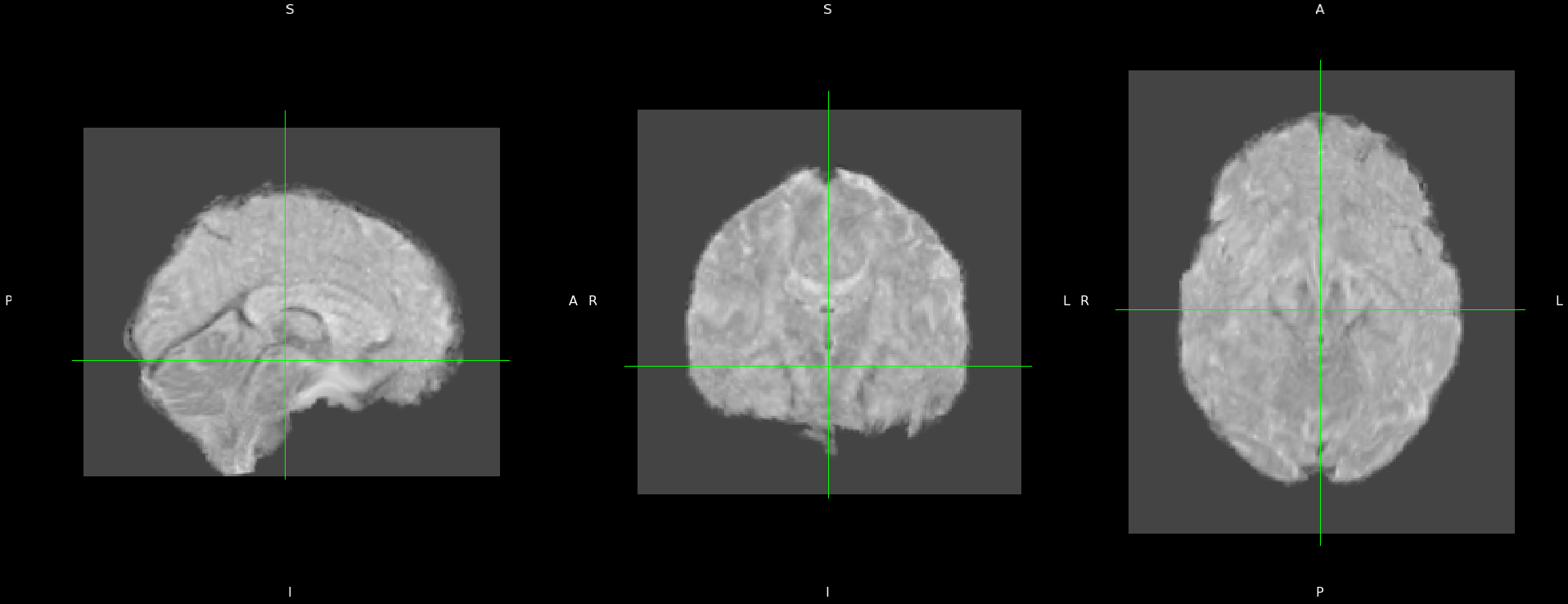

Fig 15. AD map of patient sample

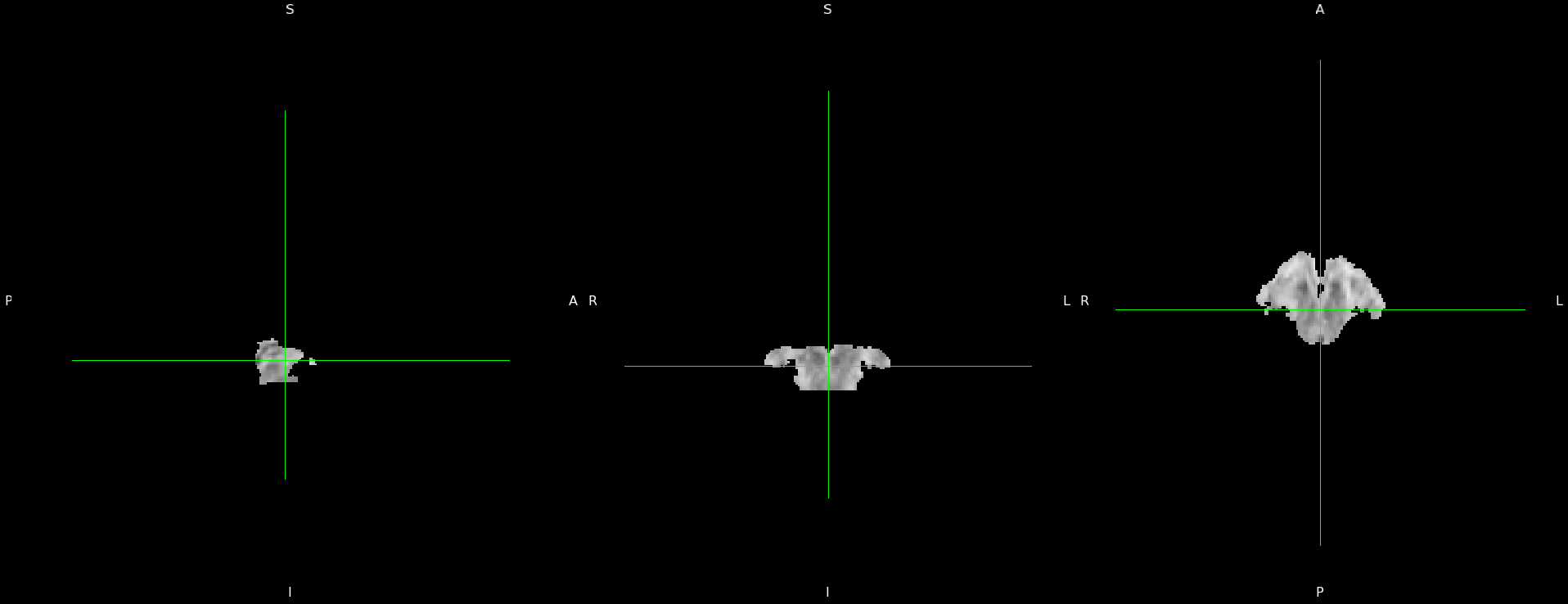

Fig 16. Midbrain segmentation of AD map

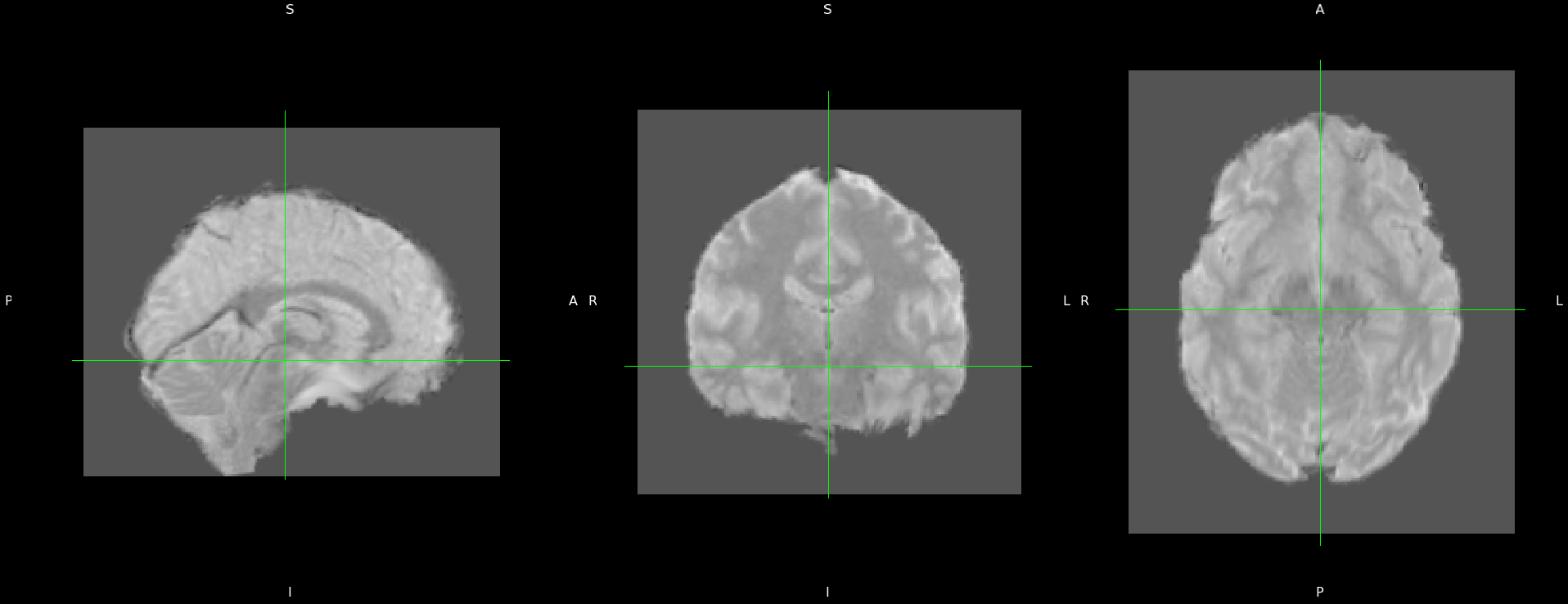

Fig 17. MD map of patient sample

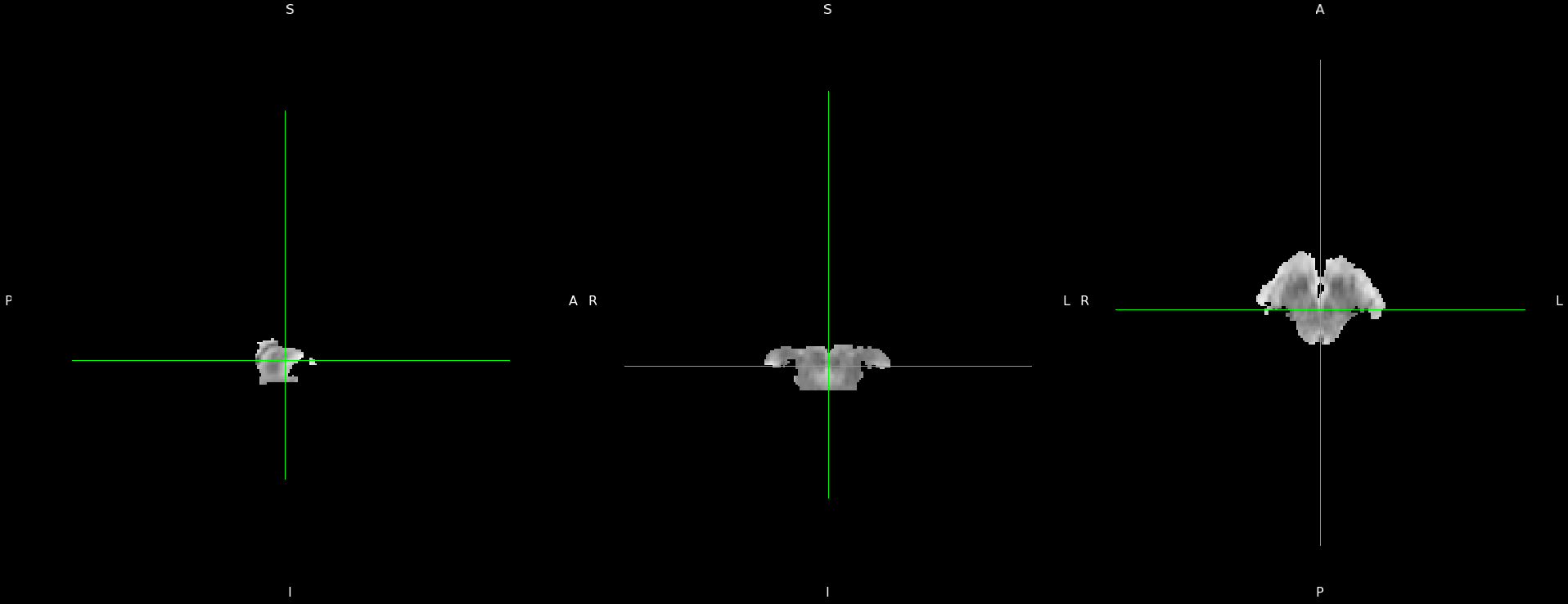

Fig 18. Midbrain segmentation of MD map

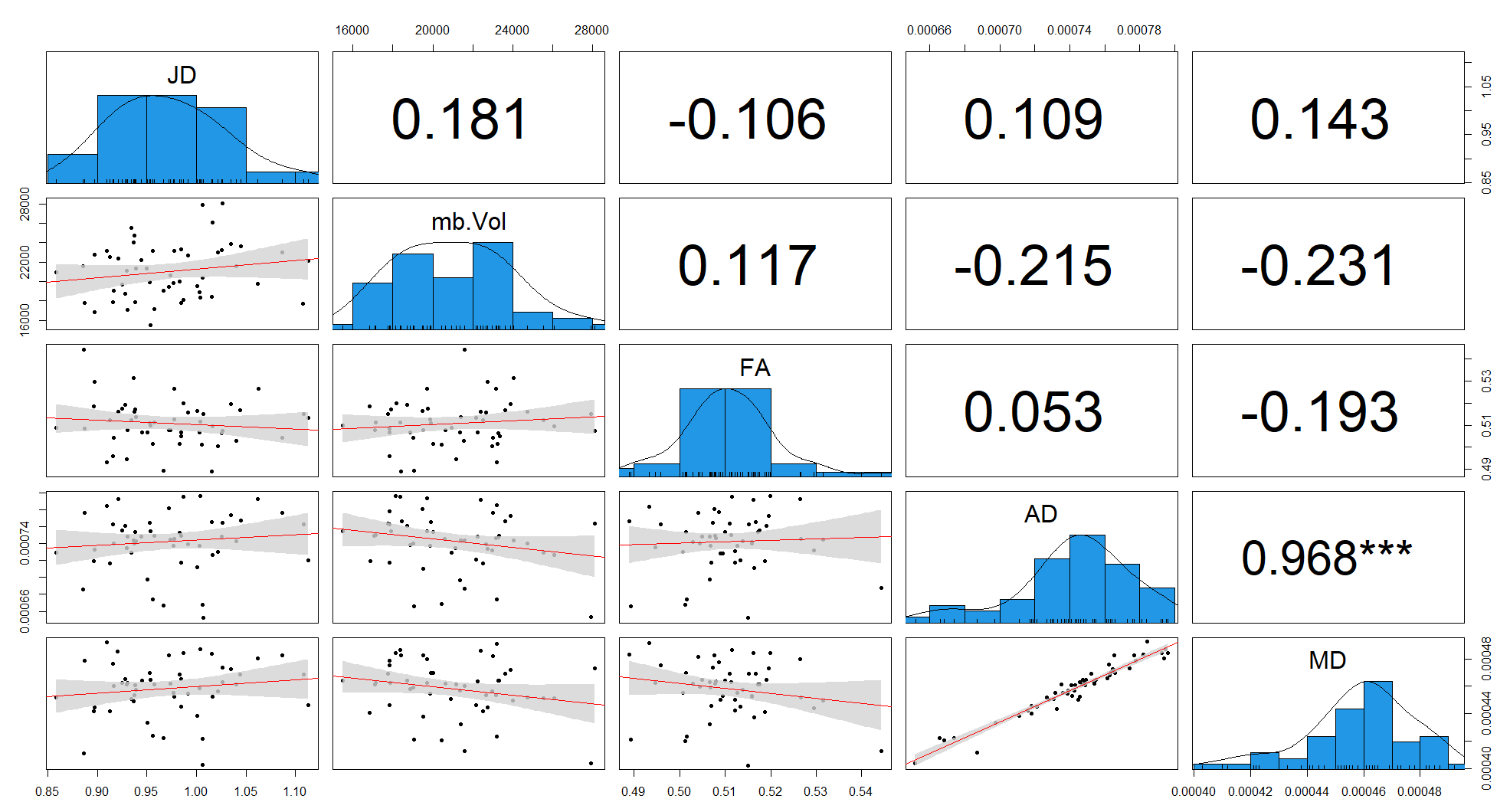

Fig 19. Correlation and distribution of MRI derived metrics in the control population (Jacobian Determinant, midbrain volumes, and DTI indices of midbrain microstructure). The diagonal shows the distribution of each score. Bottom quadrant shows the scatter plots with fitted correlation, and top quadrant shows r^2^ of the correlation (* = p < 0.05, ** = p < 0.01, and *** = p < 0.005).
